# Tmem119-EGFP and Tmem119-CreERT2 transgenic mice for labeling and manipulating microglia

**DOI:** 10.1101/624825

**Authors:** Tobias Kaiser, Guoping Feng

## Abstract

Microglia are specialized brain-resident macrophages with important functions in health and disease. To improve our understanding of these cells, the research community needs genetic tools to identify and control them in a manner that distinguishes them from closely related cell-types. We have targeted the recently discovered microglia-specific *Tmem119* gene to generate knock-in mice expressing EGFP (JAX#031823) or CreERT2 (JAX#031820) for the identification and manipulation of microglia, respectively. Genetic characterization of the locus and qPCR-based analysis demonstrate correct positioning of the transgenes and intact expression of endogenous *Tmem119* in the knock-in mouse models. Immunofluorescence analysis further shows that parenchymal microglia, but not other brain macrophages, are completely and faithfully labeled in the EGFP-line at different time points of development. Flow cytometry indicates highly selective expression of EGFP in CD11b^+^CD45lo microglia. Similarly, immunofluorescence and flow cytometry analyses using a Cre-dependent reporter mouse line demonstrate activity of CreERT2 primarily in microglia upon tamoxifen administration with the caveat of activity in leptomeningeal cells. Finally, flow cytometric analyses reveal absence of EGFP expression and minimal activity of CreERT2 in blood monocytes of the *Tmem119-EGFP* and *Tmem119-CreERT2* lines, respectively. These new transgenic lines extend the microglia toolbox by providing the currently most specific genetic labeling and control over these cells in the myeloid compartment of mice.

**Visual abstract:** 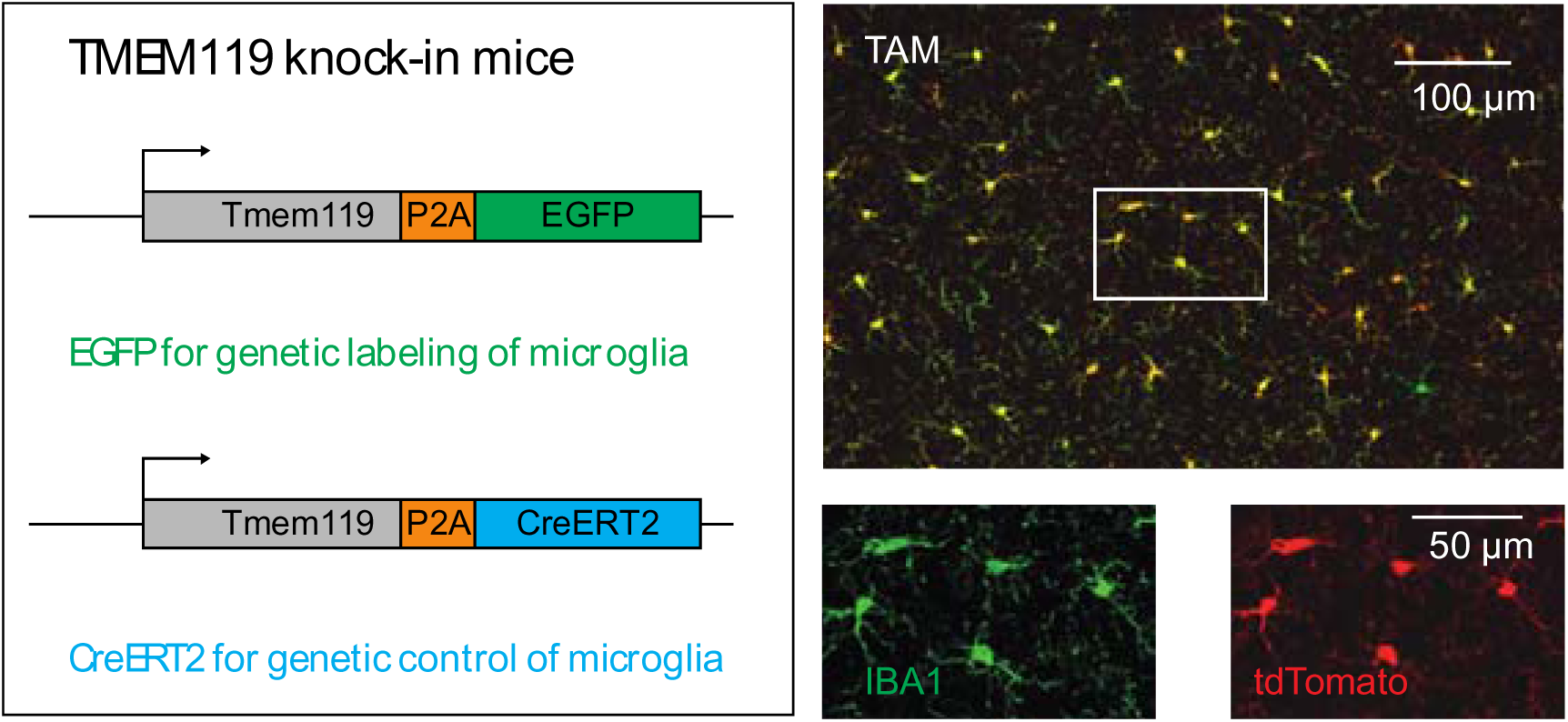

**Significance statement:** Tools that specifically label and manipulate only microglia are currently unavailable, but are critically needed to further our understanding of this cell type. Complementing and significantly extending recently introduced microglia-specific immunostaining methods that have quickly become a new standard in the field, we generated two mouse lines that label and control gene expression in microglia with high specificity and made them publicly available. Using these readily accessible mice, the research community will be able to study microglia biology with improved specificity.

## Introduction

Microglia are specialized brain-resident macrophages that comprise 5 to 12% of the glial cells in the adult mouse brain (Lawson et al., 1990). Under physiological conditions, phagocytic and secretory activity of these glia support neurogenesis, development of neuronal connectivity, and survival of neurons (Stevens et al., 2007; Sierra et al., 2010; Paolicelli et al., 2011; Schafer et al., 2012; Ueno et al., 2013; Weinhard et al., 2018). Complementing these homeostatic functions, microglia react to perturbations, which has been shown in the context of vascular injury, multiple sclerosis lesions, and neurodegeneration (Itagaki et al., 1989; Davalos et al., 2005; Ransohoff, 2016; Aguzzi and Zhu, 2017; Keren-Shaul et al., 2017; Mathys et al., 2017; O’Loughlin et al., 2018). In the future, many of these processes and their potential for therapeutic targeting will be further examined, and understanding of both homeostatic and disease-related contributions of microglia will critically depend on means to specifically identify and control them. Several mouse lines are currently available for either fluorescent labeling or Cre-expression that harness the loci of putative microglia signature genes *Tie2, Runx1, Csf1r, Aif1, Lyz2, Itgam, Sall1*, or *CX3cr1* (Clausen et al., 1999; Sasmono et al., 2003; Ferron and Vacher, 2005; Hirasawa et al., 2005; Samokhvalov et al., 2007; Ginhoux et al., 2010; Parkhurst et al., 2013; Yona et al., 2013; Buttgereit et al., 2016). Collectively, these mouse lines have been instrumental in gaining insights on microglia. However, furthering our understanding of microglia using the currently available lines is complicated by the fact that it is difficult to distinguish microglia from other closely related peripheral and central cell types such as blood monocytes as well as perivascular, choroid plexus, and meningeal macrophages (Wieghofer and Prinz, 2016; Haimon et al., 2018).

Recent advances in RNA sequencing and other cell profiling technologies have enabled the discovery of cell type-specific signature genes (Butovsky et al., 2013). Amongst these, transmembrane protein 119 (*Tmem119*) is highly and exclusively expressed in microglia in the brains of mice and humans (Bennett et al., 2016; Satoh et al., 2016). While TMEM119-specific antibodies are already widely used as a gold standard for immunohistochemical methods, faithful fluorescent reporter lines and inducible Cre-lines that allow *in vivo* observation and manipulation are currently not available. Here, we report the generation and characterization of knock-in mouse lines *Tmem119-EGFP* (JAX#031823) and *Tmem119-CreERT2* (JAX#031820), where microglia express EGFP and CreERT2, respectively, while preserving endogenous *Tmem119* expression. We demonstrate that EGFP is expressed throughout the brain and that the tag is confined to microglia only, without significantly labeling other brain macrophages. We further provide evidence that the inducible Cre is primarily active in microglia by crossing to the conditional fluorescent reporter mouse line Ai14. In these mice, we also observe activity in leptomeningeal cells that line the surface of the brain and penetrate deep into the brain ensheathing some large blood vessels. Finally, we demonstrate minimal to absent transgene expression in monocytes of the mice. These publicly available mouse lines provide valuable tools for the functional study of bona fide microglia.

## Materials and methods

### Animal work

All animal procedures were performed in accordance with the [Author University] animal care committee’s regulations. Overall, more than 30 animals of more than 10 litters were generated per line and mice were crossed to C57BL/6J up to generation N3. For all experiments, at least three independent mice were analyzed, which included both sexes and no apparent sex differences were observed.

### Generation of transgenic animals using CRISPR/Cas9

To generate *Tmem119* knock-in mice, donor DNA templates encoding ribosome-skipping peptide *porcine teschovirus-1 polyprotein* and *EGFP* (*P2A-EGFP*) or *CreERT2* (*P2A-CreERT2*) were synthesized. These sequences were flanked by sequences of 55-300 bp for EGFP and 1.5 kb for CreERT2 homologous to 5’ and 3’ regions around the *Tmem119* stop codon. These templates were injected into fertilized mouse oocytes together with a single crRNA (AGUCUCCCCCAGUGUCUAAC, Synthego) that cuts at the stop codon.

Donor DNA templates were generated by digesting pAAV-P2A-EGFP (sequence below) with XbaI and EcoRI (New England Biolabs) and inserting three gblocks (LHA, P2A-EGFP or P2A-CreERT2, RHA, purchased from IDT, sequences below) using Gibson cloning (HIFI assembly mix, New England Biolabs) according to the manufacturers protocol. Resulting plasmids were purified and sequenced. For CreERT2, the highly purified dsDNA plasmid was directly used as donor DNA in injections. For EGFP, single strand DNA (ssDNA) was produced using PCR with forward primers for left homology arms between 55 and 300 bp (5’-A*G*C*AACTGGTCCTCCTGAAA-3’ and 5’-CAAAGCCTGTGAAGGGTGGG-3’, respectively, * denoting phosphorothioate) and a reverse primer for a 55 bp right homology arm (5’-CAAAGAGGTGACCCTCAAGG-3’, with 5’ phosphorylation for lambda digest of antisense strand). Using these primers, large scale PCR with Takara PrimeStar was performed to obtain 40-60 µg of product. The PCR product was highly purified with the Qiagen PCR purification kit and subject to digestion of the antisense strand. Lambda exonuclease (New England Biolabs) was used to digest 20 µg dsDNA at 37 °C for 60 min. Complete digestion of dsDNA was confirmed by agarose gel electrophoresis and Sanger sequencing with sense- and antisense-binding primers.

Mixtures for injection of zygotes were prepared freshly on the morning of the day of injection. Briefly, water to a final volume of 100 µl was mixed with final concentrations of 10 mM TrisHcl buffer, 0.61 µM crRNA, 0.61 µM tracrRNA (sequence proprietary to Synthego) and heated to 95 °C for 5 min. Heated mixtures were cooled to room temperature for 10 min and 30 ng/µl EnGen Cas9 protein (New England Biolabs) was added. Mixtures were incubated at 37 °C for 15 min to form Cas9 to crRNA-tracrRNA complexes. Final concentrations of 5 ng/µl donor DNA (P2A-EGFP ssDNA or P2A-CreERT2 plasmid dsDNA) and 10 ng/µl recombinant RAD51 protein were added. Mixtures were kept on ice until use, when they were incubated at 37 °C for 15 min followed by centrifugation at 10.000 rpm for 1 min to prevent clogging of the micropipette.

Injection of mixtures was carried out using standardized protocols of the transgenics facility in zygotes obtained from pure C57BL/B6N mice (Taconic).

### Genetic analysis and genotyping of Tmem119 mice and Ai14

Founder mice were genetically examined by amplifying sequences spanning the 5’ and 3’ junction and including the entire inserted transgene. Specifically, high quality DNA was obtained from ear punches using a tissue DNA extraction kit according to the manufacturer’s instructions (Macherey-Nagel). One amplicon spanning approximately 1.5 kb (open arrows, Figure 1) of these sequences was generated using Q5 polymerase with primers 5’ofLHA-F (GCCTCTGTCACTTAAGTTGG) and P2A-R (GCTTCAGCAGGCTGAAGTTA). A second amplicon of about 2.5-3.5 kb (closed arrows, Figure 1) was generated using primers Tmem-LHA-P2A-F (CAGTGTCGGAAGCGGAGCTA) and 3’ofRHA-R (GAAAGAGGAAGCTAGAAGGG). Both amplicons were purified and then sequenced using primers spanning the entire length using Sanger sequencing (Genewiz). Trace files were aligned to *in silico* assemblies and analyzed using Snapgene.

**Figure 1.**
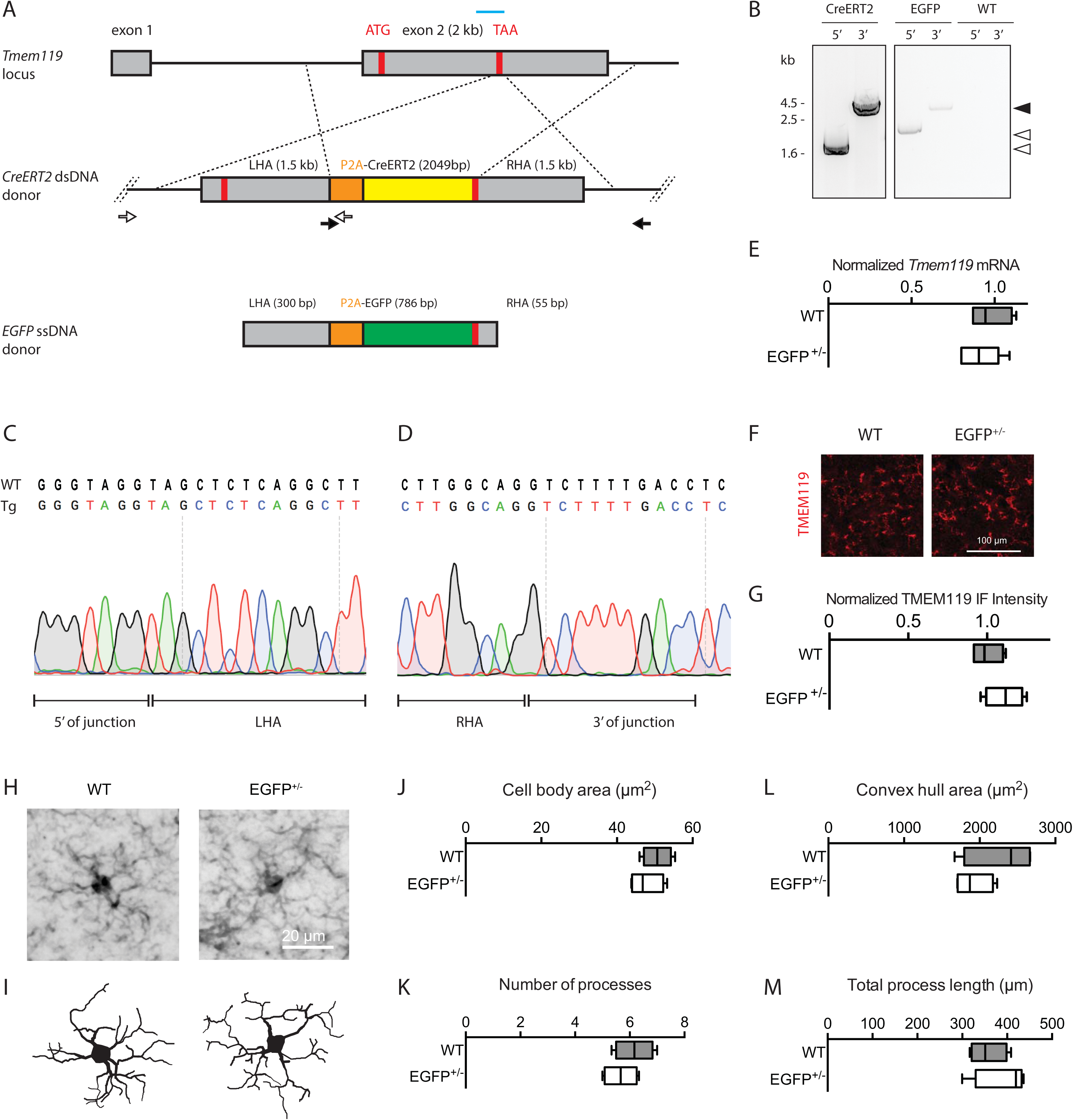
Generation and validation of *Tmem119-EGFP and Tmem119-CreERT2* knock-in lines. **A**, Experimental strategy using a guide RNA (blue bar) to introduce a double-strand break at the *Tmem119* stop codon in mouse zygotes and an injection mixture containing *EGFP* ssDNA template with short homology arms of 55-300 bp or *CreERT2* dsDNA template with long 1.5 kb homology arms. Each knock-in is designed to cause in-frame insertion of P2A peptide followed by *EGFP* or *CreERT2* and a stop codon. Not drawn to scale. **B**, Confirmation of insertion by amplification of fragments spanning the 5’ and 3’ junctions using sets of primers binding inside the template and outside the homology arms (open and closed arrows in A with arrowheads indicating corresponding products). **C and D**, Sanger sequencing across the 5’ and 3’ junction of the CreERT2 line and comparison to wildtype (WT) sequence indicate seamless insertion in the transgenic mice (Tg). **E**, qPCR analysis of gene expression for *Tmem119* in wildtype controls and *Tmem119-EGFP^+/−^* knock-in animals. N=4 WT, N=6 *Tmem119-EGFP^+/−^* mice. **F**, Representative TMEM119 immunoreactivity in wildtype controls and *Tmem119-EGFP^+/−^* knock-in animals. **G**, Relative intensity of TMEM119 immunostaining signal. N=4 WT, N=4 *Tmem119-EGFP^+/−^* mice. **H and I**, Representative IBA-immunostained microglia and corresponding neurolucida traces in WT and *Tmem119-EGFP^+/−^* knock-in mice. **J-M**, Quantified cell body area, number of processes, convex hull area, and total process length of microglia in WT and *Tmem119-EGFP^+/−^* knock-in mice. N= 12 microglia from 4 mice per genotype. ssDNA, single strand DNA; dsDNA, double strand DNA; TAA (red bar), stop codon; Kb, kilo bases.

To genotype mice, ear tissue was prepared using an alkaline buffer (25 mM NaOH, 0.2 mM Na_2_EDTA, pH=12) at 95 °C for 30 min. The lysed tissue solution was neutralized using an acidic buffer (40 mM TrisHcl, pH=5). PCR using primers WT-F GTCAGGAGGAGGCCCAGGAA, EGFP-F CTGCTGCCCGACAACCACTA, CreERT2-F ACCGCCTACATGCGCCCACT, and Common-R GTTTCCTGGGGTGCACCAGA yielded products of 400 bp for the wildtype allele and 320 bp for the EGFP or CreERT2 allele.

Ai14(RCL-tdT) mice were obtained from JAX (Stock 007914) and maintained in house. Homozygous Ai14 mice were bred to heterozygous *Tmem119-CreERT2* mice to generate offspring for the analysis. Ai14 mice were genotyped according to the protocol provided by JAX.

### RNA isolation and quantitative PCR (qPCR)

Brain hemispheres from adult *Tmem119-EGFP^+/−^* mice were dissected and snap-frozen in liquid nitrogen. Highly pure RNA was isolated using an RNeasy purification kit (Qiagen) following the manufacturer’s instructions. Equal amounts of RNA (1.5 µg per sample) were reverse transcribed using an iScript Advanced cDNA synthesis kit (Bio-Rad). Resulting cDNA was diluted 1:50 in ultrapure water. qPCR was carried out using SsOAdvanced Universal SYBR Green Supermix (Bio-Rad) on a CFX96 realtime system. Primers used were exon-spanning whenever possible and of the following sequences: Tmem119-F CCTTCACCCAGAGCTGGTTC, Tmem119-R GGCTACATCCTCCAGGAAGG, GAPDH-F GCCTTCCGTGTTCCTACC, GAPDH-R CCTCAGTGTAGCCCAAGATG, b-actin-F CTAAGGCCAACCGTGAAAAG, b-actin-R ACCAGAGGCATACAGGGACA. Differential gene expression analysis was performed using built-in software for the Bio-Rad CFX96 realtime system.

### Tamoxifen administration

Tamoxifen (Sigma, T5648) was dissolved in corn oil at 20 mg/ml under agitation for several hours in the dark and kept at room temperature for 2–3 days. Using needles from Harvard Apparatus (52-4025), separately housed adult animals were fed via oral gavage for 3 days. Needles were rinsed between days and syringes were replaced. At time of administration, the mice weighted approximately 19-26 g and received 250-400 µl of the 20 mg/ml tamoxifen solution, corresponding to 0.2 mg per g body weight. Postnatal day 2 mice weighing approximately 2 g received 5 µl of the 20 mg/ml solution, corresponding to 50 µg per g body weight per day for three consecutive days. The health of the mice was monitored daily. At least 7 days week after the last dose, mice were sacrificed by transcardiac perfusion.

### Immunofluorescence staining and imaging

Adult mice were deeply anesthetized and perfused with 25 ml phosphate buffered saline (PBS) followed by 25 ml 4% paraformaldehyde (PFA) in PBS. Early postnatal mice (P3) were perfused with 8 ml PBS and 8 ml PFA. Brains were surgically removed and postfixed in the fixative at 4 °C for 24 h. Fixed brain were washed in PBS once and sliced into 100 µm-thick sagittal slices using a Leica VT1000S. Slices were washed twice in PBS, permeabilized in 1.2% TX100 in PBS for 15 min, washed twice in PBS, and subject to incubation in blocking solution (5% normal goat serum, 2% bovine serum albumin, 0.2% TX100 in PBS). Blocked sections were incubated with primary antibodies for IBA1 (1:500, Synaptic Systems, 234006), GFP (1:1000, Invitrogen, A11122, 1:500, Aves Labs, GFP-1020), TMEM119 (1:1000, Abcam, ab209064), Olig2 (1:1000, Millipore, AB9610), NeuN (1:1000, Millipore, MAB377), GFAP (1:1000, Sigma, G9269), or CD163 (1:200, Abcam, ab182422, antigen retrieval required) for 24 h at 4 °C. Primary antibody incubation was followed by three washes in PBS and incubation with species-matched and Alexa fluorophore-conjugated secondary antibodies raised in goat (Invitrogen, 1:1000) for 2 h. DAPI (1:10.000) was included in a washing step or secondary antibody incubation. Slices were washed three times in PBS and mounted and coverslipped using vectashield H-1000 mounting medium.

For immunostaining of CD163, an antigen retrieval step was included. Briefly, vibratome slices were washed twice in PBS, incubated in a retrieval buffer (10 mM Sodium citrate, pH 8.5) for 5 min, followed by incubation in the retrieval buffer at 80 °C for 30 min. Sections were cooled to room temperature and washed twice in PBS. Blocking and immunostaining were then carried out as described above.

For imaging, slides were scanned on an Olympus Fluoview FV1000 fixed stage confocal microscope (high power magnifications) or Olympus BX61 epifluorescence microscope (sagittal section montage) using built-in software. For co-expression analysis in the *Tmem119-EGFP* line, maximum projections of 4 z-stacks from three 20X high-power fields (approximately 100 cells per region) were counted manually.

For microglia morphology analysis, z-stacks were acquired at 20X magnification with 2X digital zoom and 1.2 µm step size. Images were imported as stacks into Neurolucida software. Cell bodies in the middle of the stacks with complete process arbors were selected and manually traced. Raw trace files were imported into Neurolucida explorer software and convex hull area, number of processes and total process length were determined using the convex hull analysis and branched structure analysis functions. Rendered traces were exported as monochrome vector graphics. Cell body areas were separately determined in FIJI using the polygon tool.

### Cell suspension preparation

Microglia and blood monocytes for single cell suspensions for flow cytometry were prepared as follows. Mice were deeply anesthetized with isofluorane and transcardially perfused with ice-cold HBSS. During perfusion, 2 ml of whole blood was collected from the right atrium into tubes containing 40 µl of 10% (w/v) EDTA. The 2 ml of whole blood were added to 40 ml of red blood cell lysis buffer (Abcam, ab204733) and incubated for 10 min at room temperature to lyse red blood cells. The suspension containing lysed RBCs and white blood cells was spun down at 300 g for 5 min at 4 °C, resuspended in 10 ml ice-cold HBSS, and pelleted again at 300 g for 5 min at 4 °C. WBCs were resuspended in 500 µl ice-cold FACS buffer (HBSS, 0.5% BSA, 1 mM EDTA) and subject to staining. In parallel with the WBC enrichment, brains were rapidly dissected into 2 ml ice-cold HBSS and cerebella and brainstem were removed. Brains were minced into small pieces and transferred to a dounce-homogenizer containing 5 ml ice-cold HBSS with 20 µg/ml DNase I (Worthington, DPRF, LS006343). Tissue chunks were homogenized with 15 loose and 15 more tight strokes and the homogenate was transferred to a 50 ml falcon tube through a pre-wet 70 µm strainer. The strainer was rinsed with HBSS to top off the volume of each sample to 10 ml. Filtered homogenates were transferred to 15 ml falcon tubes and spun at 300 g for 5 min at 4 °C. Supernatants were carefully removed, and pellets resuspended in 10 ml of ice-cold 40% Percoll in HBSS. Samples were spun at 500 g for 30 min at 4 °C with full acceleration and deceleration. Myelin and debris from the supernatant were carefully removed and pellets resuspended in 10 ml ice-cold HBSS. Following another spin at 300 g for 5 min at 4 °C, the supernatant was removed and the microglial pellet resuspended in 1 ml FACS buffer (0.5% BSA HBSS, 1 mM EDTA).

### Staining for flow cytometry

Suspensions of white blood cells and microglia were transferred to 2 ml Eppendorf microcentrifuge tubes, and a small fraction of sample was removed for single-color controls. To stain dead cells, live/dead violet (1:500 to 1:1000, Thermofisher, L34955) was added and incubated for 5 min on ice. Tubes with live/dead-stained cells were topped off to 2 ml with FACS buffer (0.5% BSA HBSS, 1 mM EDTA) and spun down at 300 g for 5 min at 4 °C. Pellets were resuspended and incubated with 1:200 Mouse Fc Block (BD, 2.4G2, 553142) on ice for 15 min. Samples were incubated with 1:200 rat anti CD45-APC/Cy7 (Biolegend, 103115) and rat anti Cd11b-PE/Cy5 (Biolegend, 101209) at 4 °C. Tubes were topped off to 2 ml with ice-cold FACS buffer and microglia pelleted at 300 g for 5 min at 4 °C. Supernatants were removed and microglia resuspended in 500 µl. Resuspended microglia were filtered through corning strainer polystyrene tubes (Corning, 352235). Flow cytometry data was acquired on Aria II, Fortessa HTS, and LSRII HTS, and analyzed using Flowjo.

### Statistical analysis

Quantitative data from the qPCR, immunofluorescence, and microglia morphology experiments were analyzed using GraphPad prism. Testing for normality was not possible due to N=4-6 (four to six independent mice were used per group). Not assuming Gaussian distribution of the data, a nonparametric test (Mann-Whitney) was used to compare median expression in both groups.

### DNA sequcnes pAAV-P2A-EGFP

#### pAAV-P2A-EGFP

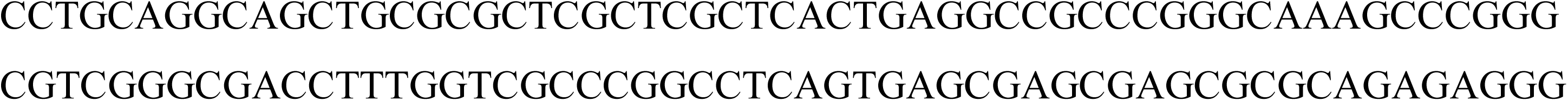

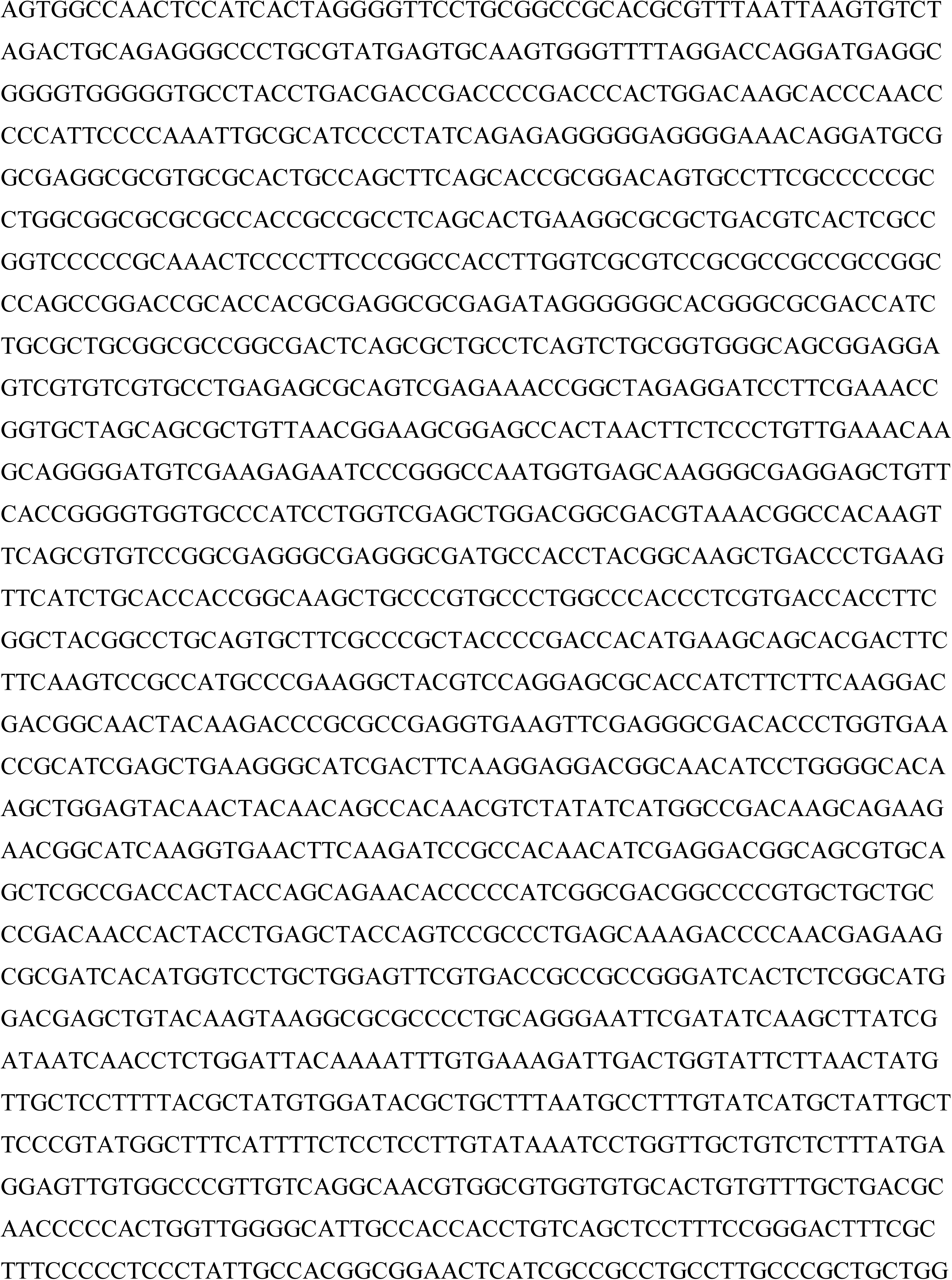

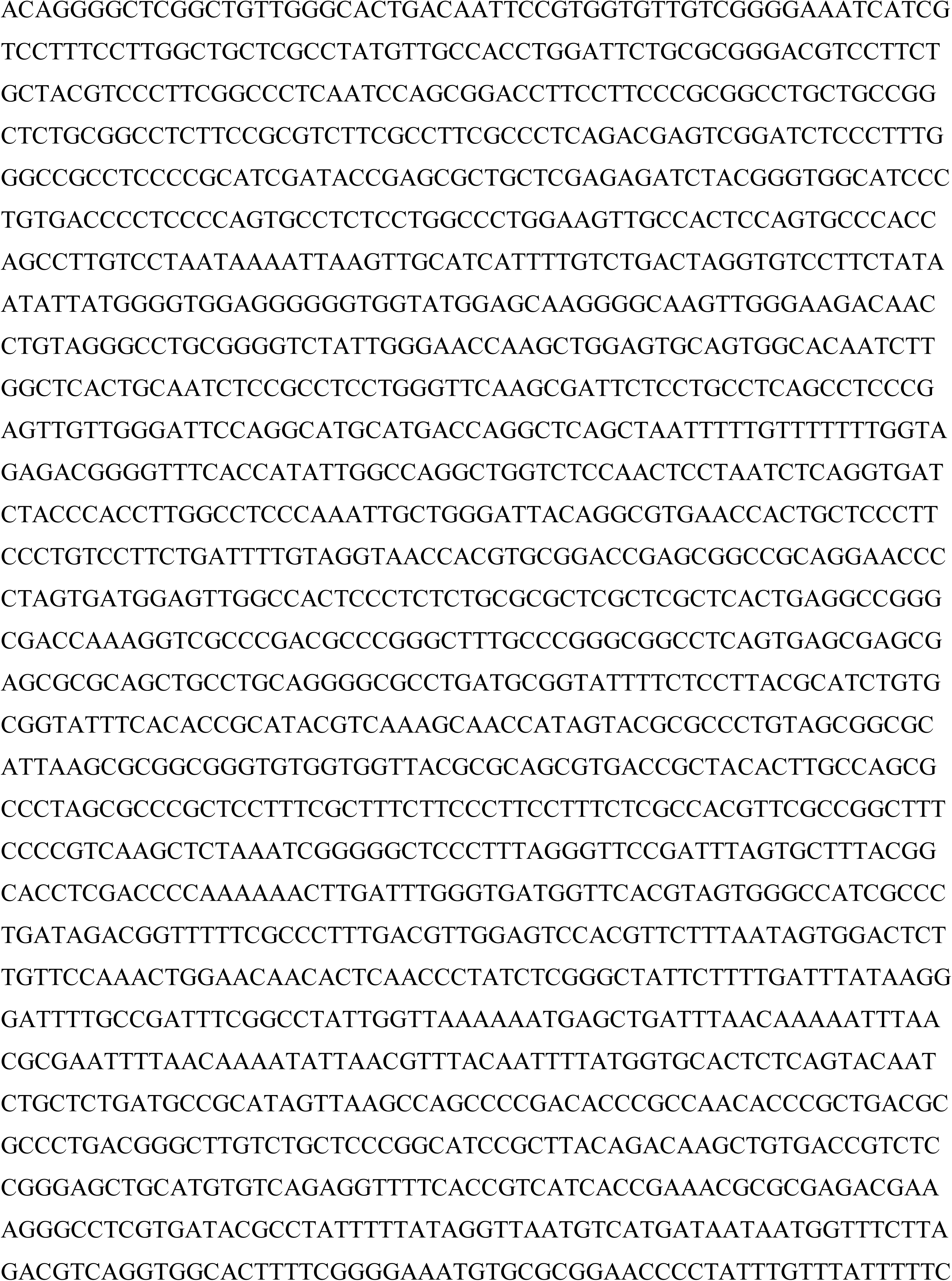

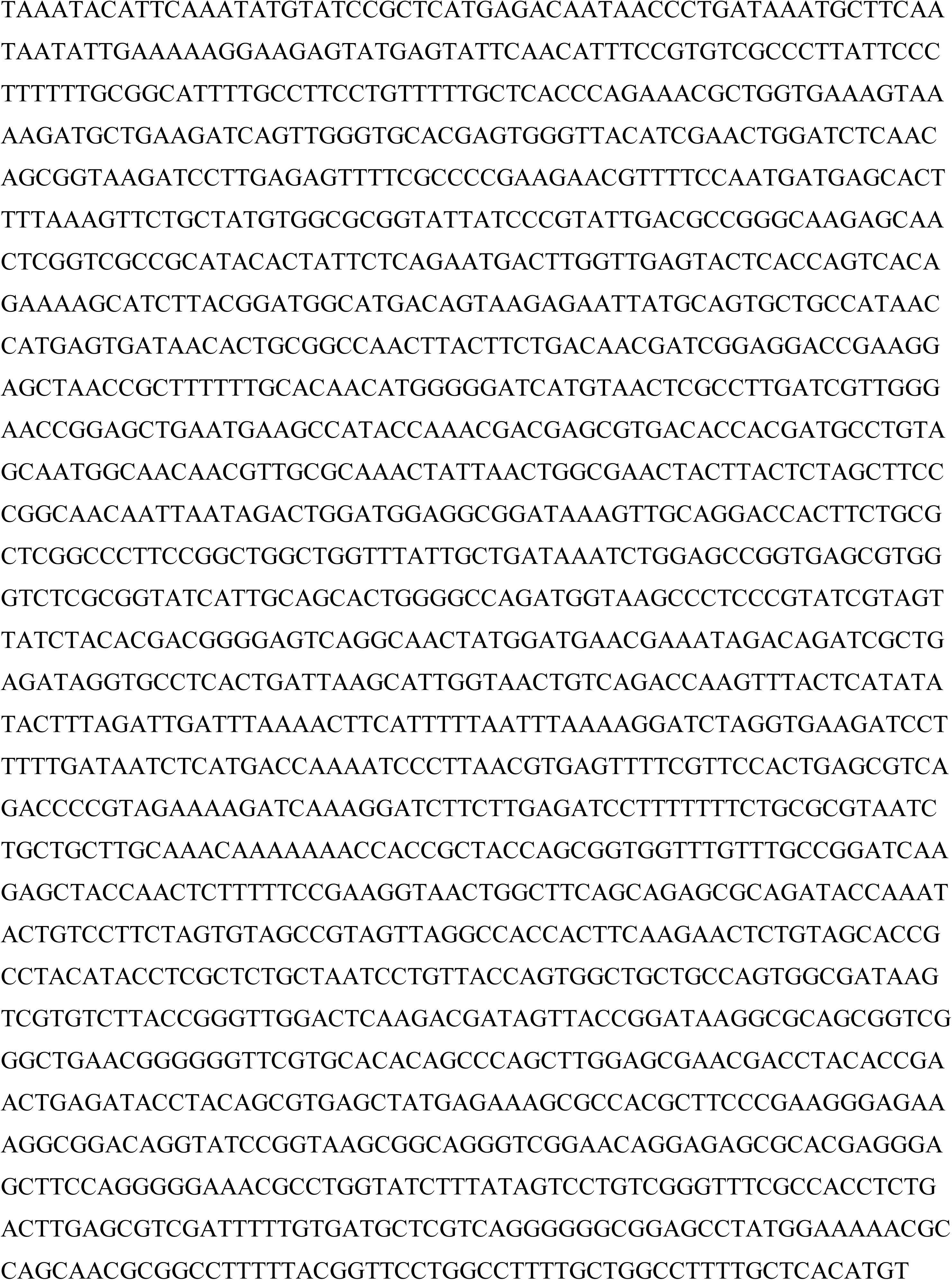

### LHA

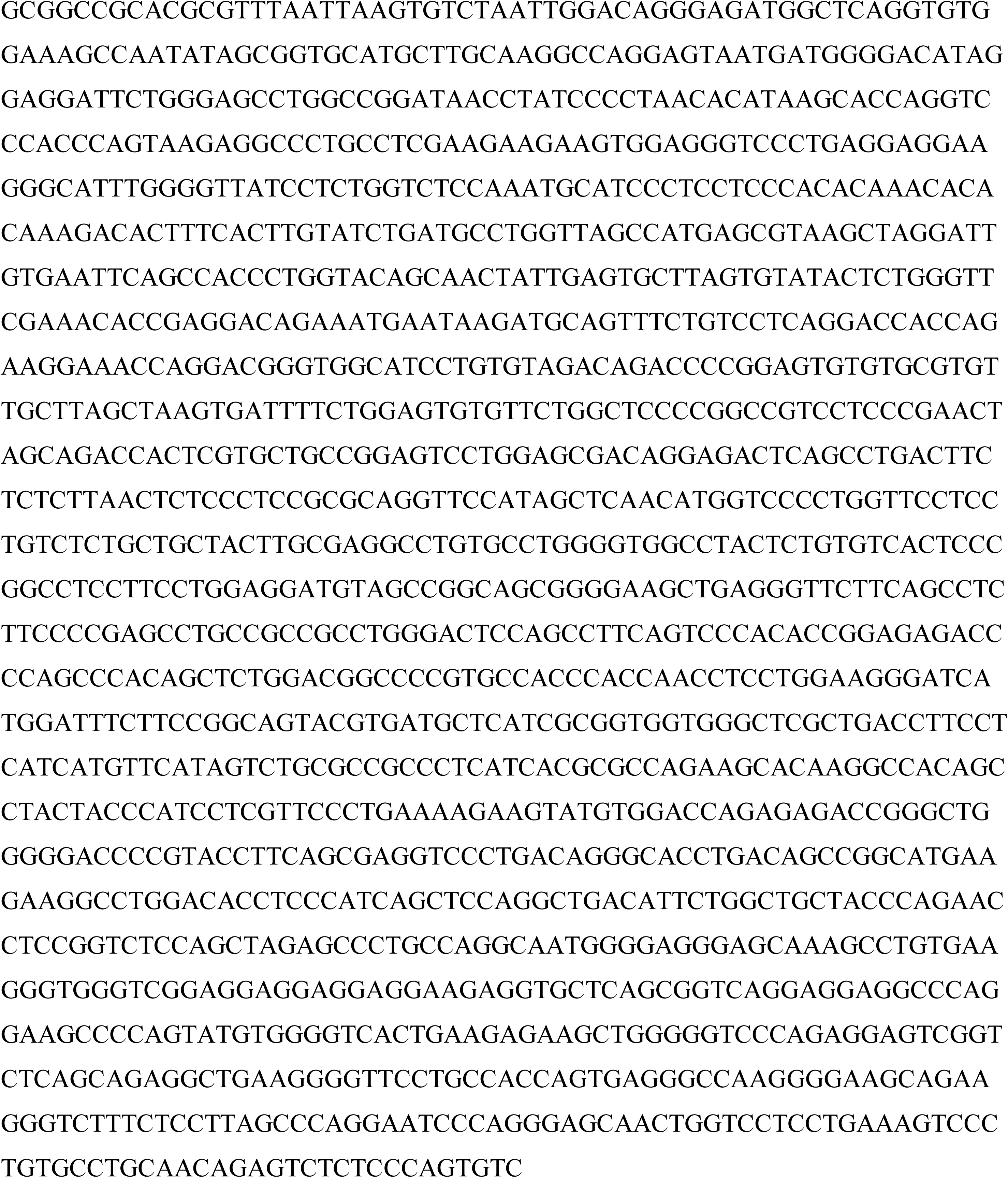

### P2A-CreERT2

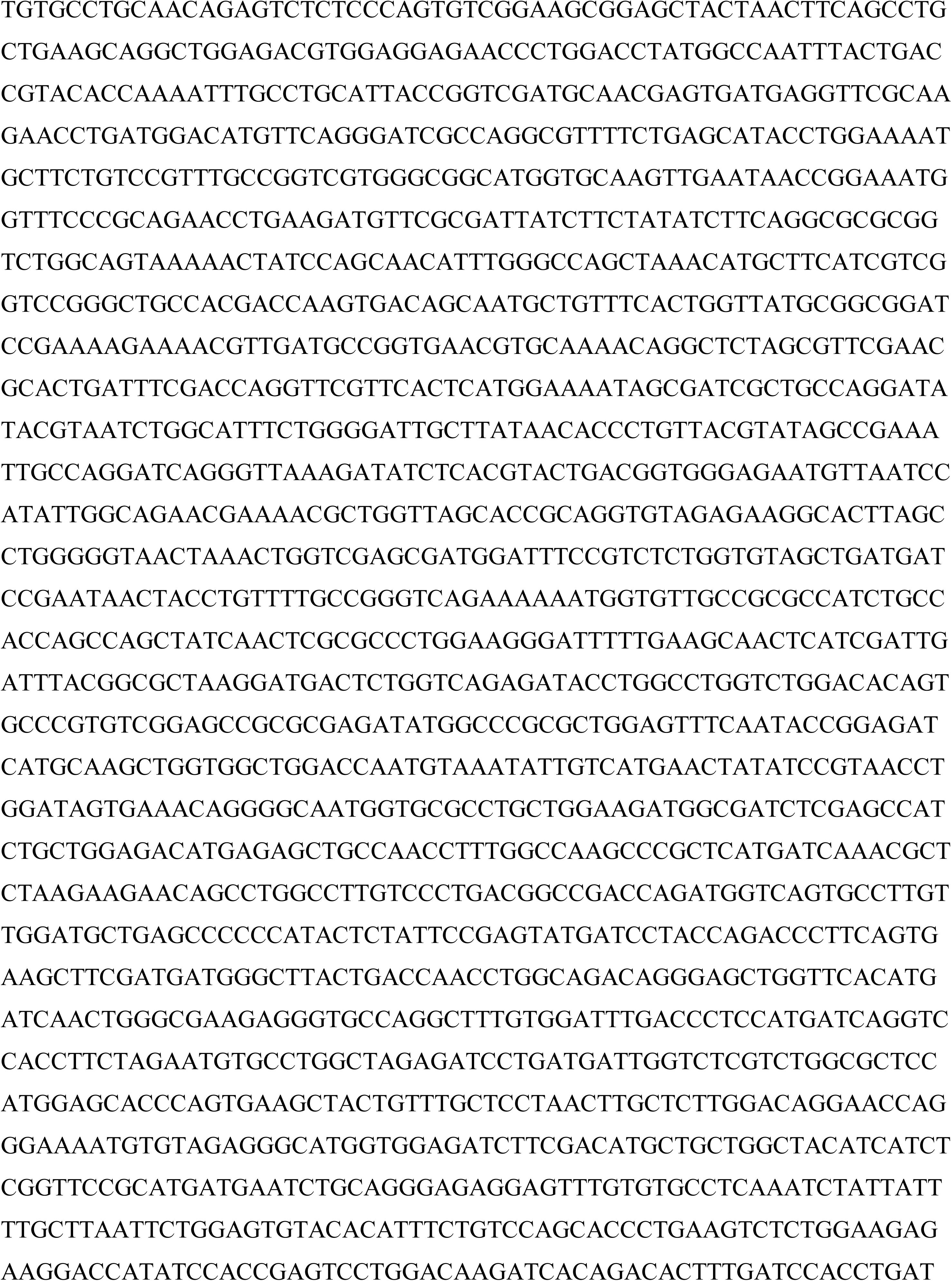

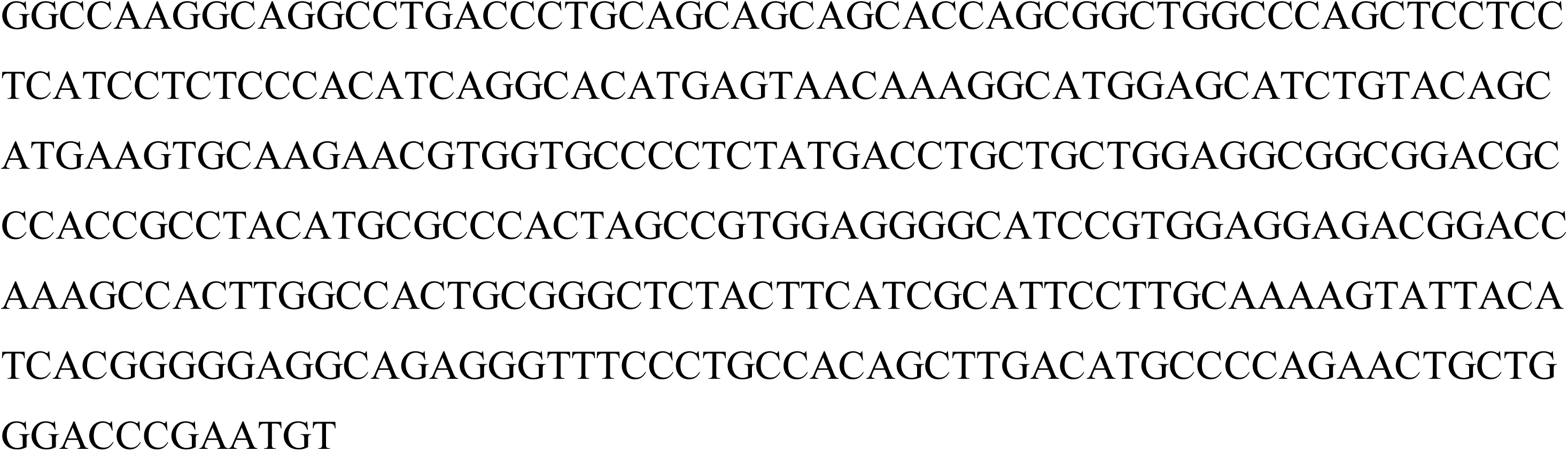

### P2A-EGFP

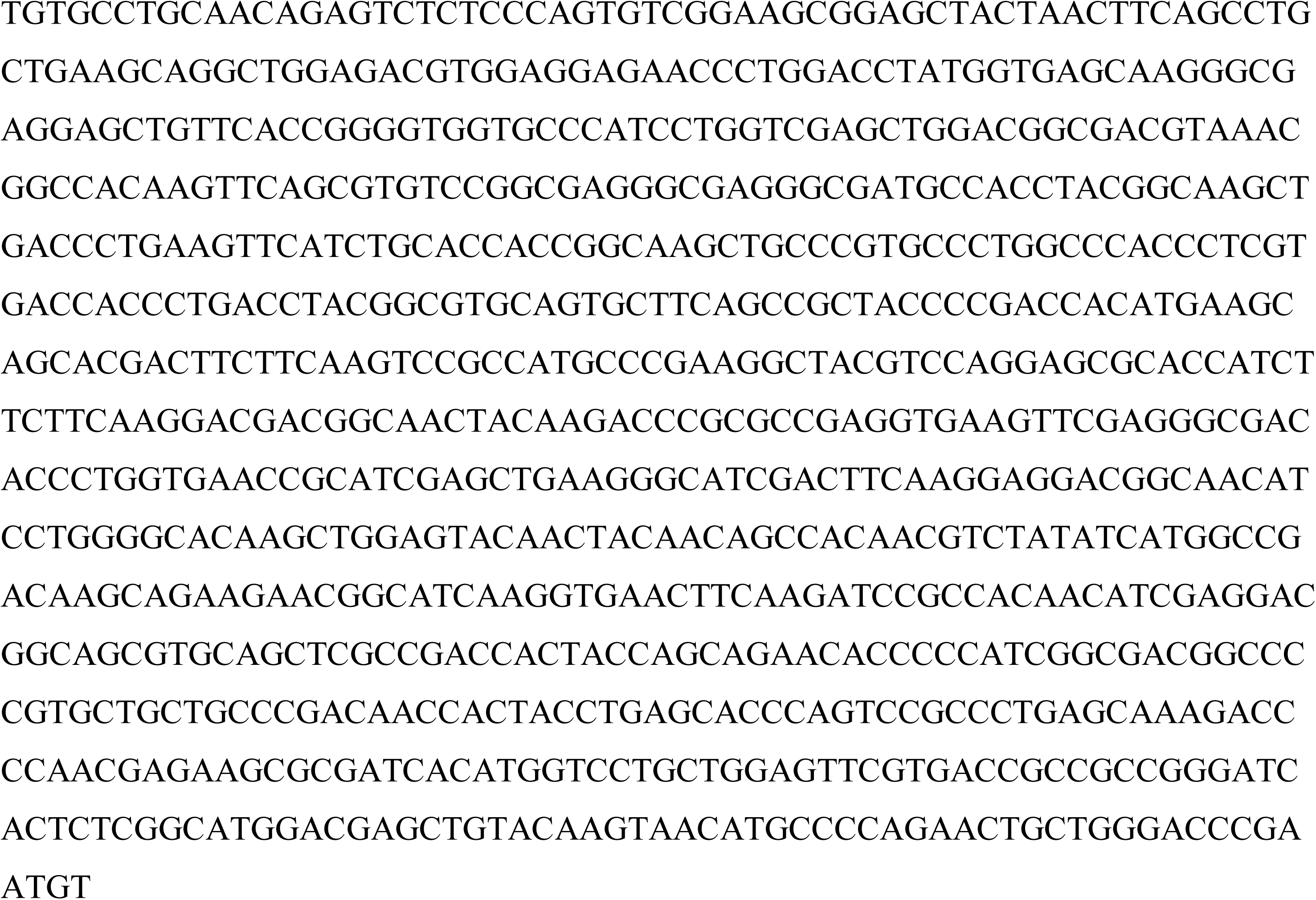

### RHA

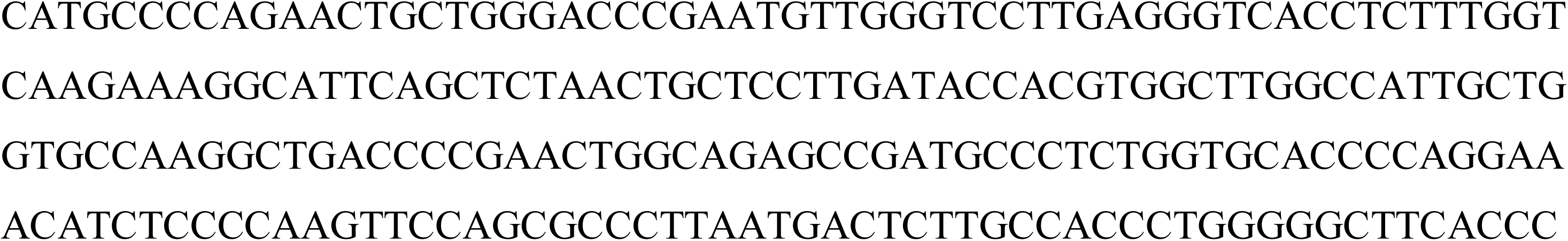

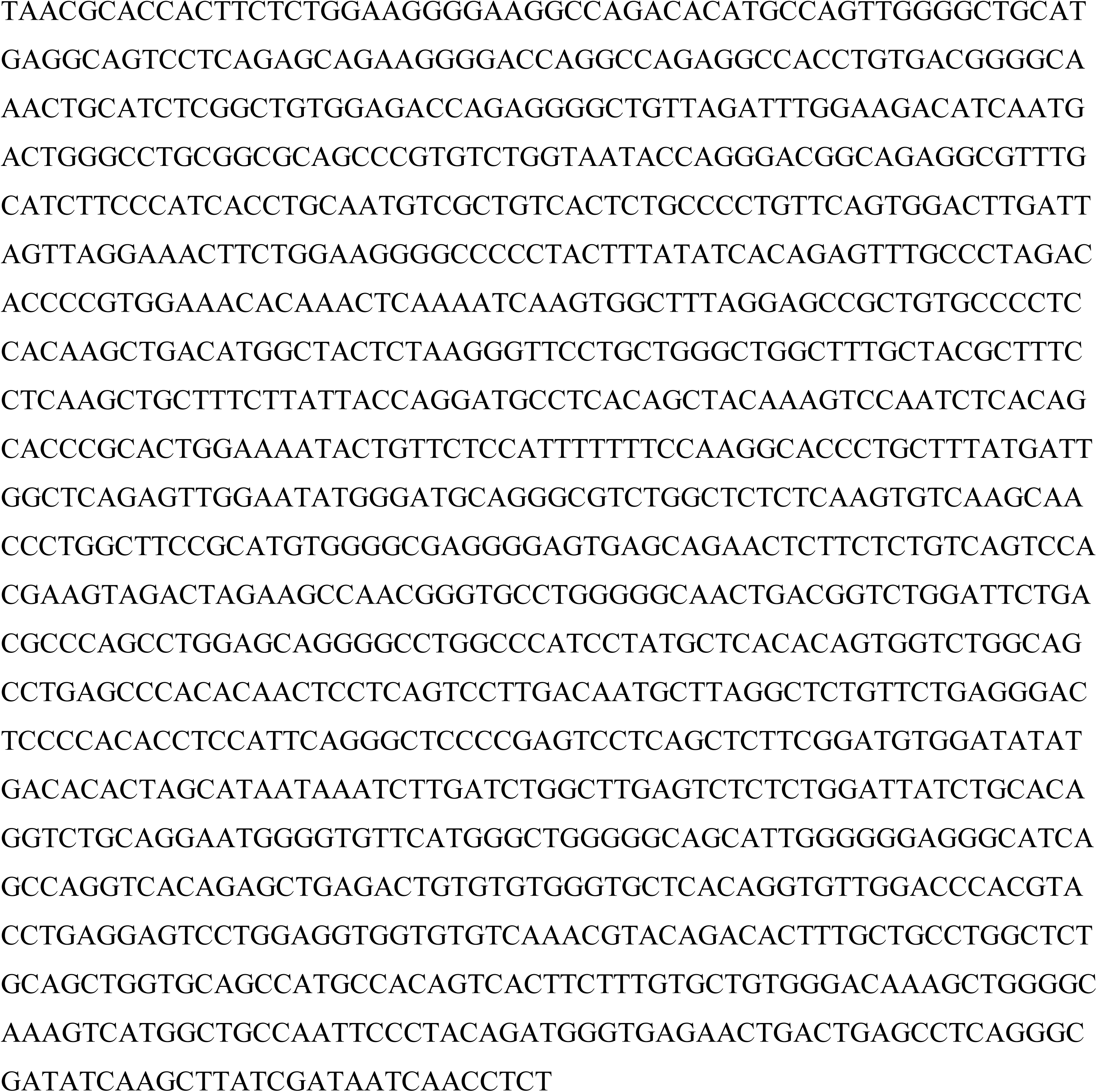

## Results

### Generation and genetic confirmation of Tmem119-EGFP and Tmem119-CreERT2 knock-in lines

Access to highly specific fluorescent reporter lines for imaging and Cre-driver lines for functional manipulation is critical in our quest for understanding the roles of microglia in health and disease. Previously generated lines efficiently target microglia, but also other cells of the monophagocytic system (Wieghofer and Prinz, 2016). Amongst a group of more recently discovered microglia-specific genes, we chose targeting the Tmem119 gene to generate such mouse lines (Bennett et al., 2016). Specifically, we opted for a transgenic approach that harnesses the Tmem119 locus to drive expression, while preserving endogenous Tmem119 expression. We designed a CRISPR/Cas9 strategy to insert EGFP or CreERT2, preceded by ribosome-skipping peptide porcine teschovirus-1 polyprotein (P2A), into the stop codon of murine *Tmem119* (Figure 1A). Two founder lines were established for Tmem119-EGFP and 5 for Tmem119-CreERT2.

To determine whether the donor DNA was inserted correctly, a PCR-based approach was used with primers spanning each junction. For both of the lines generated, specific 5’ and 3’ junction PCR products were obtained, while no bands were observed with DNA from wildtype animals (Figure 1B). This indicates specific insertion of the transgenes into the *Tmem119* locus. To further examine scarless integration at the junctions, PCR products from the CreERT2 line were subject to Sanger sequencing covering the junctions. Both 5’ and 3’ sequencing traces show endogenous bases corresponding to the WT sequence in intron 1 and the 3’ UTR (Figure 1C, D). Additional sequencing confirmed absence of mutations across the entire targeted allele in both lines. Together, these results confirm the generation of *Tmem119-EGFP* and *Tmem119-CreERT2* lines.

While the function of endogenous *Tmem119* is currently unknown, both its temporally distinct pattern and high expression level suggest functional importance of this gene (Bennett et al., 2016). We thus designed a knock-in approach with the objective of leaving endogenous expression intact. To test whether *Tmem119* expression is actually unaffected, we quantified *Tmem119* transcript levels in the *Tmem119-EGFP* line by qPCR. Examination of *Tmem119* expression in WT and *Tmem119-EGFP*^+/−^ mice revealed that transcript levels are comparable between genotypes (Figure 1E, WT median 0.9717, n=4, EGFP^+/−^ median 0.9306, n=6, p= 0.5667^a^). We further examined TMEM119 protein expression using immunofluorescence and found no difference between wildtype and *Tmem119-EGFP^+/−^* knock-in mice (Figure 1F-G, WT median 0.9835, n=4, EGFP^+/−^ median 1.117, n=4, p=0.20^b^) Together, these data support the conclusion that *Tmem119* knock-in mice were generated with precision.

To further test whether the genetic engineering could affect the properties of microglia, we examined basic morphological features of reconstructed immunostained microglia in wildtype and *Tmem119-EGFP^+/−^* knock-in mice (Figure 1H-I). Using this approach, we determined that the cell body area (Figure 1J, WT median 50.67 µm^2^, EGFP^+/−^ median 46.82 µm^2^, n=12 microglia from 4 mice per genotype, p=0.3429^c^), number of processes (Figure 1K, WT median 6.167, EGFP^+/−^ median 5.667, n=12 microglia from 4 mice per genotype, p=0.4571^d^), convex hull area (Figure 1L, WT median 2413 µm^2^, EGFP^+/−^ median 1871 µm^2^, n=12 microglia from 4 mice per genotype, p=0.4857^e^), and total process length (Figure 1M, WT median 351.2 µm, EGFP^+/−^ median 419.6 µm, n=12 microglia from 4 mice per genotype, p=0.3428^f^) are comparable between microglia from control and knock-in mice. These data suggest that the genetic modification does not affect basic microglial properties.

### EGFP is faithfully expressed in microglia

Microglia populate the parenchyma in virtually all regions across the brain. To determine the extent of EGFP expression in the newly generated Tmem119-EGFP line, we used confocal microscopy on brain slices. Native EGFP fluorescence and antibody-enhanced signal were visible across the brain in sagittal brain sections prepared from postnatal day 25 (P25) *Tmem119-EGFP*+/- mice (Figure 2A). To further assess whether EGFP is expressed in microglia, immunofluorescence labeling of the endogenous microglia proteins IBA1 and TMEM119 was employed. High power confocal microscopy shows that EGFP-positive cells in the parenchyma are IBA1 and TMEM119-positive (Figure 2B-E). Completion and fidelity of expression were further assessed quantitatively in several brain regions. We found that 96.1 ± 2.0 % of the EGFP-expressing cells in the somatosensory cortex are IBA-1 expressing microglia and 98.2 ± 3.3 % of IBA1-positive cells in this region are labeled by EGFP (Table 1). In the striatum, 99.2 ± 2.1 % of the cells labeled with EGFP were IBA1-expressing and 97.2 ± 1.8 % of IBA1-positive cells displayed EGFP fluorescence. Labeling of microglia in the thalamus and hippocampus was 98.5 ± 0.8 % and 98.9 ± 1.9 % complete and 96.9 ± 2.3 % and 98.5 ± 3.5 % specific, respectively. To further confirm EGFP expression in microglia across the brain, we used the *Tmem119-EGFP* line in flow cytometry (Figure 2F-N). We gated for single, live cells (Figure 2F-I) and detected EGFP^+^ cells among CD45^+^ cells (Figure J-M). We determined that all CD11b^+^CD45lo microglia (Figure 2K) expressed EGFP (Figure 2N). In addition, to more comprehensively assess specificity of EGFP expression, we immunostained oligodendrocytes, neurons, and astrocytes. We found no evidence for expression of EGFP in these major CNS celltypes (Figure 2O-Q). Together, these data indicate highly complete and specific labeling of parenchymal microglia in the *Tmem119-EGFP* reporter line.

**Table 1:** Quantification of EGFP labeled microglia in Tmem119-EGFP mice.

| Brain Region | Completeness<br>(EGFP<sup>+</sup>Iba<sup>+</sup>/Iba<sup>+</sup>) | Fidelity<br>(EGFP<sup>+</sup>Iba<sup>+</sup>/EGFP<sup>+</sup>) |
| --- | --- | --- |
| Somatosensory cortex | 98.2 ± 3.3 % | 96.1 ± 2.0 % |
| Striatum | 97.2 ± 1.8 % | 99.2 ± 2.1 % |
| Thalamus | 96.9 ± 2.3 % | 98.5 ± 0.8 % |
| Hippocampus | 98.5 ± 3.5 % | 98.9 ± 1.9 % |

**Figure 2.**
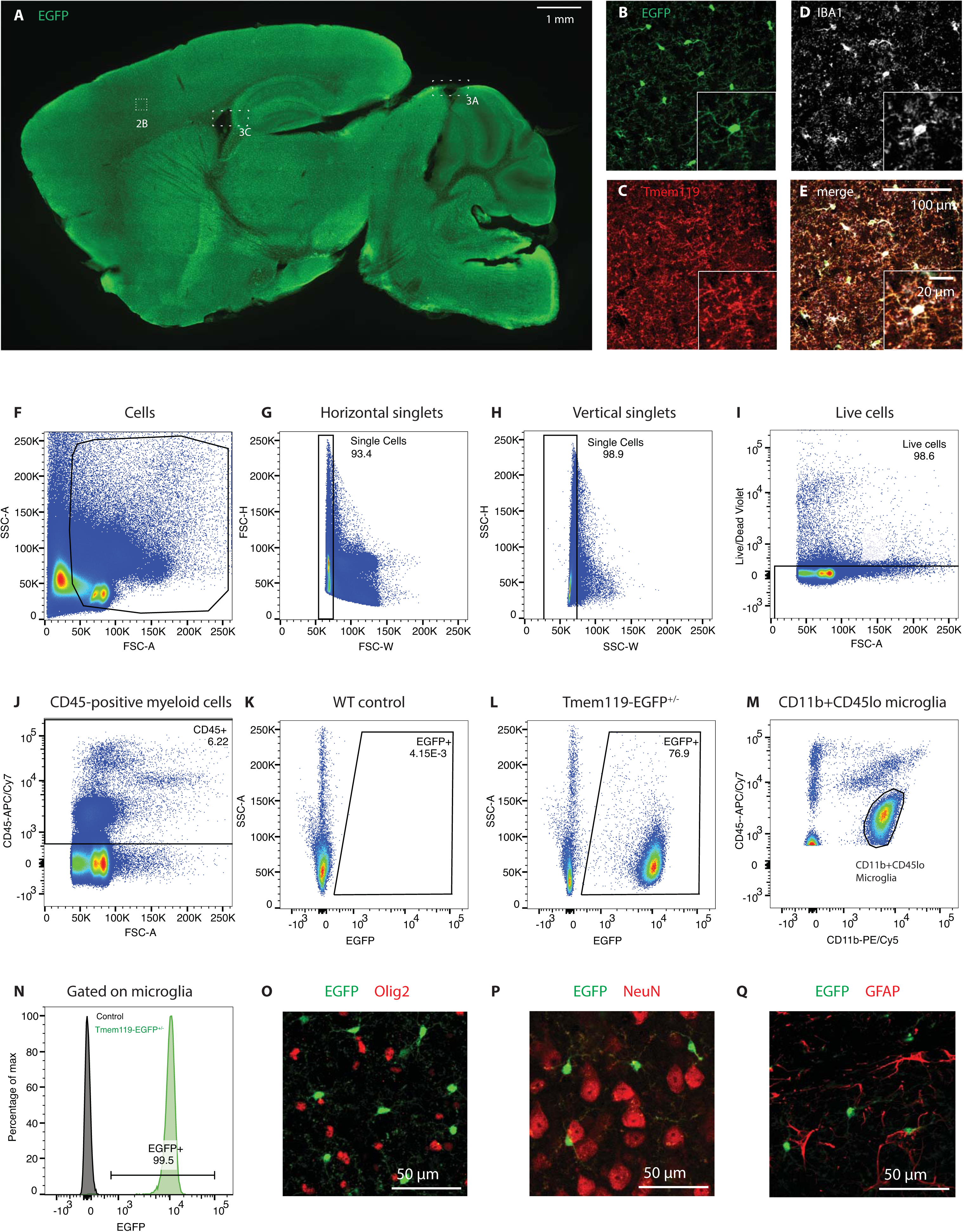
EGFP labels microglia in all regions across the brain. **A**, Representative epifluorescence image of a sagittal section of a P25 *Tmem119-EGFP^+/−^* mouse stained with an anti-GFP antibody for improved signal-to-noise ratio (data for one of three mice shown). **B-E**, High-power confocal micrographs of native EGFP fluorescence (B, green) in TMEM119 (C, red) and IBA1 (D, white) microglia in the cortical region outlined in A. FIJI-calculated composite image showing all three labels (E). Boxes 3A and 3B cross-reference areas shown in figure 3.**F-J**, Flow cytometry analysis showing gating for of single, live, CD45-positive cells (one of three independent experiments shown). Numbers in or adjacent to outlined areas indicate percent cells in each gate. **K and L**, Representative density blots showing EGFP expression in WT and *Tmem119-EGFP^+/−^* mice (pre-gated on CD45^+^ cells). **M**, Representative density plot showing CD11b^+^CD45lo cells corresponding to microglia (pre-gated on CD45^+^ cells). **N**, Histogram showing fraction of CD11b^+^CD45lo microglia expressing EGFP. **O-R**, Representative immunostaining for oligodendrocytes (O, Olig2, red), neurons (P, NeuN, red), and astrocytes (Q, GFAP, red) in EGFP-expressing *Tmem119-EGFP^+/−^* mice. One of three independent experiments shown.

All microglia express IBA1, but not all IBA1-expressing cells are parenchymal macrophages. In fact, non-parenchymal macrophages such as meningeal, perivascular, and choroid plexus macrophages express the myeloid- and macrophage marker IBA1 (Prinz and Priller, 2014). To determine whether EGFP is expressed in these non-parenchymal macrophages, we examined EGFP immunofluorescence in IBA1-expressing cells in the meninges and choroid plexus, which can be readily discerned by their morphology and IBA1 immunoreactivity. High power confocal images from both regions, including adjacent parenchyma, show that, while parenchymal microglia express the EGFP label, neither IBA1-positive meningeal (Figure 3A, B) nor choroid plexus macrophages express EGFP (Figure 3C, D).

**Figure 3.**
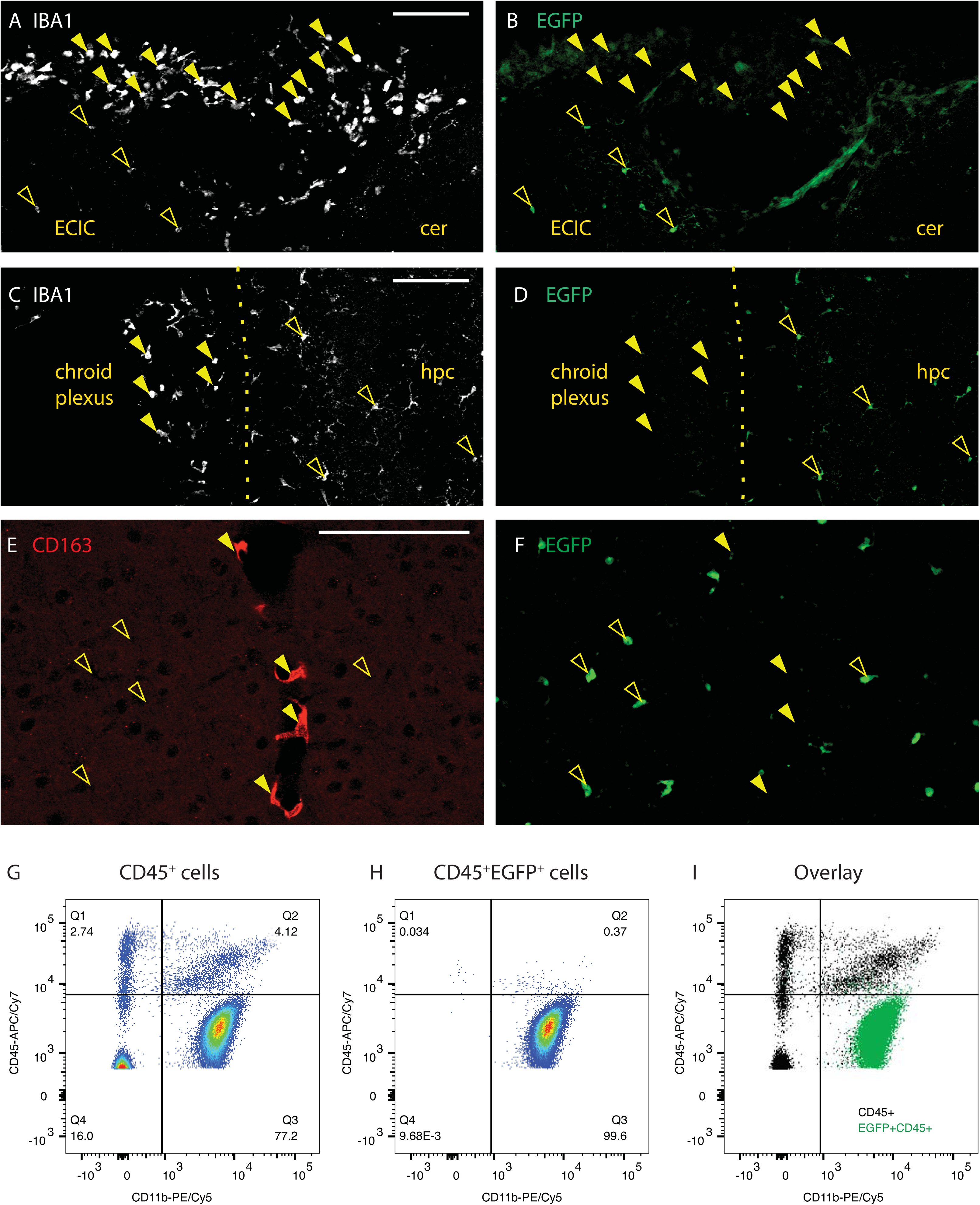
EGFP discerns parenchymal microglia from other brain macrophages. **A**, IBA1 labels meningeal macrophages and parenchymal microglia from the region indicate with a dashed box in figure 2A. **B**, EGFP labels only parenchymal macrophages (open arrowheads), but not meningeal IBA1-positive macrophages (closed arrowheads). Diffuse fluorescence is associated with pia. ECIC, external cortex of inferior colliculus; Cer, cerebellum. Scale bar 100 µm. **C and D**, Choroid plexus IBA1^+^ macrophages (closed arrowheads) from the region boxed in figure 2A are not labeled with EGFP, while adjacent parenchymal microglia are (open arrowheads). Hpc, hippocampus. Scale bar 100 µm. **E and F**, Perivascular CD163-expressing macrophages (E, red, closed arrowheads) are not labeled with EGFP (F, green, open arrowheads). Scale bar 100 µm. **G-I**, Flow cytometry analysis of CD11b and CD45 expression in pre-gated single live CD45^+^ cells and EGFP-expressing CD45^+^ cells of *Tmem119-EGFP^+/−^* mice shown as density plots (G, CD45^+^ and H CD45^+^ EGFP^+^) and overlay (I, black CD45^+^, green CD45^+^ EGFP^+^). One of three independent experiments (N ≥ 3 mice per group) shown. Numbers in or adjacent to outlined areas indicate percent cells in each gate.

In contrast to the clear anatomical location of choroid plexus and meningeal macrophages, sites of perivascular macrophages are not easily discernible anatomically. To examine whether perivascular macrophages are labeled in the *Tmem119-EGFP* line, we stained for CD163, a well-characterized marker for perivascular macrophages (Kim et al., 2006). CD163 readily identified perivascular macrophages, which did not express EGFP (Figure 3E-F). Adjacent microglia were clearly labeled with EGFP.

To further assess the specificity of EGFP expression in microglia beyond anatomical location-based immunofluorescence analysis, we used flow cytometry (Figure 3G-I). Examining CD45^+^EGFP^+^ cells, we determined that 99.6% are CD11b^+^CD45lo cells corresponding to microglia (Figure 3H). We further detected that 153 of 2212 CD11b^+^CD45int/hi cells (6.9%) corresponding to non-microglial cells express EGFP (Figure 3G-I). Together, these data indicate the capacity of the *Tmem119-EGFP* line to distinguish parenchymal microglia from other brain macrophages with good selectivity.

### EGFP is expressed in microglia of Tmem119-EGFP mice at early postnatal stages

Microglia populate the brain during early embryonic development (Ginhoux et al., 2010). *Tmem119* mRNA is expressed throughout microglial development, but antibody-based methods currently consistently label TMEM119 only as of later postnatal stages, typically postnatal day 14 (P14) and later (Bennett et al., 2016). This makes it difficult to identify and discern microglia from monocytes at earlier time points. To determine whether the *Tmem119-EGFP* line would specifically label microglia at early postnatal stages, we carried out immunofluorescence microscopy at postnatal days 1, 3, and 6. EGFP expression in IBA1-positive cells was readily observed in several regions examined, suggesting translation of *Tmem119-P2A-EGFP* transcript at this stage (Figure 4A-L). Surprisingly, we also found EGFP expression associated with blood vessels of P1 mice (Figure 4A-C), but not P3 or later, indicating that the transcript is also processed in cells of the intracerebral microvascular compartment at this stage. Together, these data indicate utility of the *Tmem119-EGFP* line for early developmental studies of microglia.

**Figure 4.**
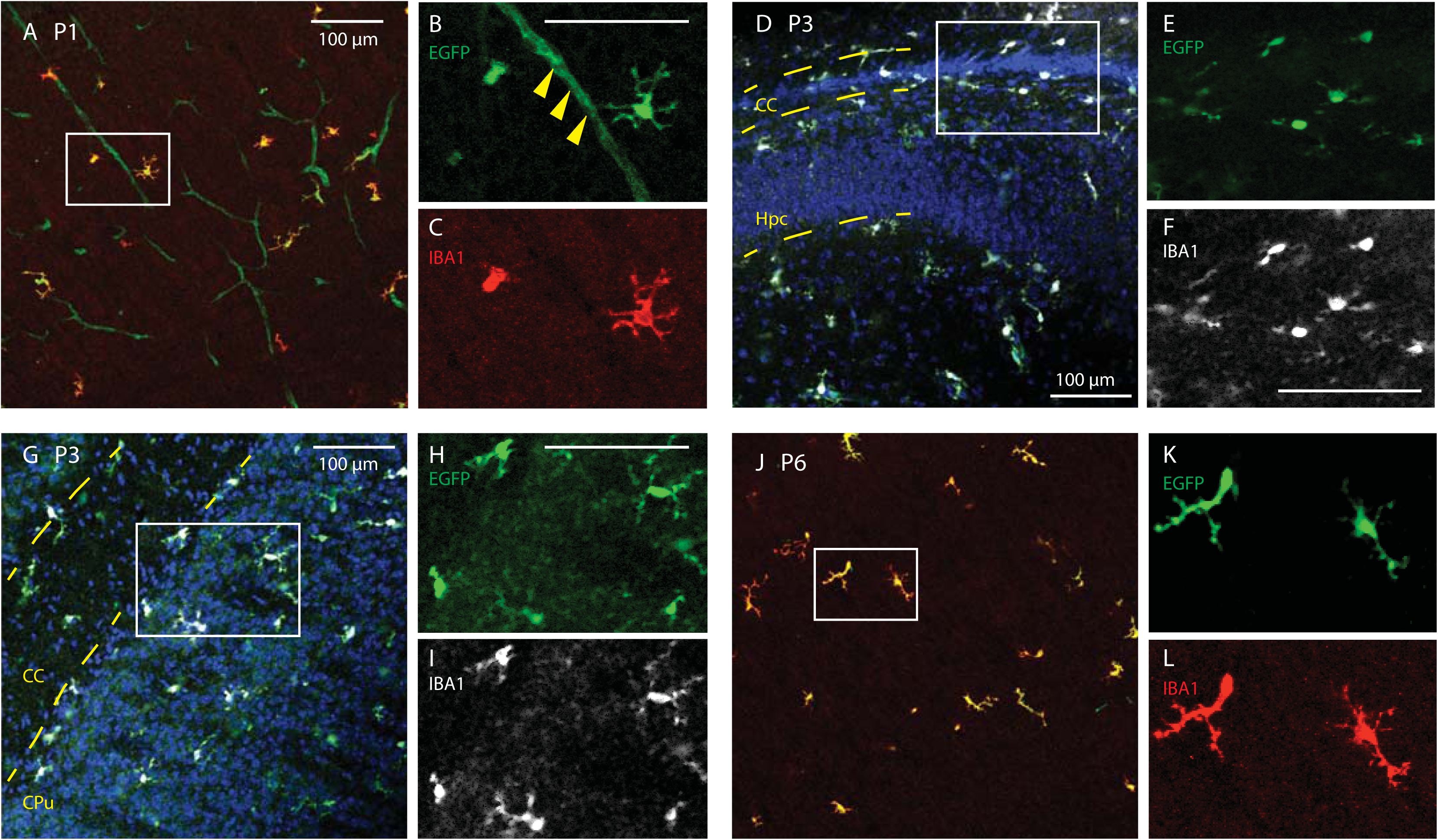
*Tmem119-EGFP* mice label microglia at early postnatal stages. Sagittal slices from postnatal day 1, 3 and 6 (P1, P3, P6) mice were stained against IBA1. **A-C**, High power confocal images of the somatosensory cortex show EGFP expression in IBA-positive cells and blood vessels at P1 (solid arrowheads in magnified panel). **D-F**, Confocal micrographs around the forceps major of corpus callosum (CC) and hippocampus (Hpc) show EGFP-labeled, IBA1-positive microglia at P3. **G-I**, Confocal micrographs of the striatum (CPu) and corpus callosum (CC) show EGFP labeling of IBA-positive cells at P3. **J-L**, Confocal micrographs of the P6 cortex reveal EGFP-expression in IBA-immunostained cells. Error bars 100 µm.

### Cre activity is specifically inducible in microglia of Tmem119-CreERT2 mice

Cre activity in CreERT2 fusion protein is induced by the estrogen receptor antagonist tamoxifen and can be assessed in the presence of a reporter construct. To assess whether *Tmem119-CreERT2* mice display Cre activity in parenchymal microglia upon induction, we crossed *Tmem119-CreERT2* mice to the Ai14 reporter line (Madisen et al., 2010), which expresses tdTomato in the presence of Cre (Figure 5A). We administered tamoxifen daily for 3, 5, or 10 days to adult mice and 3 days to postnatal day 2 (P2) pups, and compared recombination efficiency and fidelity for these different paradigms (Figure 5A). The mice were sacrificed 9-10 days after the last dose and subjected to immunofluorescence staining. High power images from confocal microscopy showed tdTomato-labeling of IBA1-expressing microglia in mice that received tamoxifen (Figure 5B-D) and absence of tdTomato expression without tamoxifen (Figure 5E-G). Confocal micrographs also indicated absence of tdTomato expression in other major cell types of the CNS, such as oligodendrocytes, neurons, and astrocytes (Figure 5H-J). Assessment of the completeness of labeling as the ratio of tdTomato^+^IBA1^+^-double positive microglia (tdT^+^IBA1^+^) to all parenchymal microglia (IBA1^+^) revealed very high efficiency of recombination (Figure 5K). Similarly, we examined the fidelity of labeling which represents the fraction of actual parenchymal microglia (tdT^+^IBA1^+^) amongst all tdTomato-labeled cells (tdT^+^) in different parenchymal regions, and found high specificity close to 100 percent (Figure 5L). Of note, increasing the days of tamoxifen administered to mice appeared to increase the fraction of microglia undergoing recombination. At the same time, prolonged administration of tamoxifen correlated with slightly reduced fidelity (Figure 5K-L). We further sought to assess the specificity of tdTomato expression using flow cytometry. Among a total of 1690 CD11b^+^CD45int/hi cells in the CNS, 33 were positive for tdTomato, indicating minimal, yet nonzero, expression in non-microglial myeloid cells.

**Figure 5.**
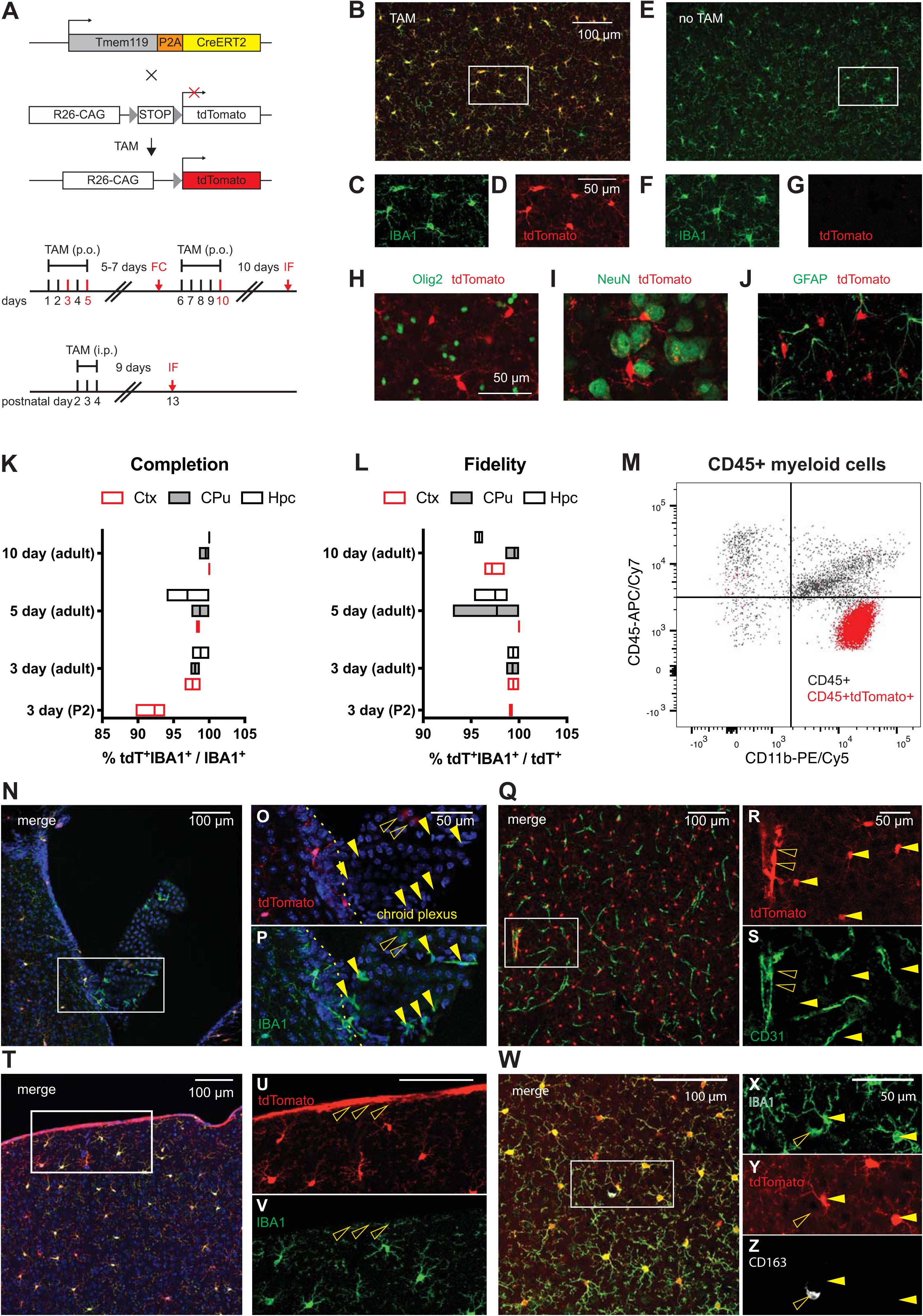
*Tmem119-CreERT2* mice effectively recombine a conditional allele in microglia. **A**, *Tmem119-CreERT2* mice were crossed to *Ai14(RCL-tdT)-D* mice and tamoxifen (TAM) was administered to induce tdTomato expression. Different sets of adult mice received tamoxifen per os (p.o) for 3, 5 or 10 days and a set of neonatal mice received TAM for 3 days intraperitoneally (i.p). The mice were sacrificed for flowcytometry (FC) 7 days after the 3-day dosing paradigm or for immunofluorescence (IF) 9 or more days after dosing. **B-D**, Representative confocal micrograph showing tdTomato expression in IBA-positive microglia upon tamoxifen administration. **E-G**, Absence of tdTomato expression in untreated animals. **H-J**, Representative immunostaining for oligodendrocytes (H, Olig2, green), neurons (I, NeuN, green), and astrocytes (J, GFAP, green) in tdTomato-expressing microglia of *Tmem119-CreERT2^+/−^; *Ai14(RCL-tdT)-D*^+/−^* mice. One of three independent experiments shown. **K and L**, Percentage completion **(K)** and fidelity **(L)** of the labeling in different brain regions for the different dosing schemes. Hpc = Hippocampus. CPu = Caudate-Putamen. Ctx = Cortex. N=2 mice for 3 days, 3 mice each for 5 and 10 days of TAM in adults, and 3 mice for 3 days administration at P2 = postnatal day 2. Floating bars = min to max, line at mean. **M**, Flow cytometry analysis of CD11b and CD45 expression in pre-gated single live CD45^+^ cells and tdTomato-expressing CD45^+^ cells of *Tmem119-CreERT2^+/−^; Ai14(RCL-tdT)-D^+/−^* mice shown as overlay (I, black CD45^+^, red CD45^+^ tdTomato^+^). One of two representative experiments (N=3 mice). **N-O**, Confocal micrograph of the choroid plexus and adjacent parenchyma. **Q-S**, Confocal micrograph of CD31-immunostained blood vessels (green) shows tdTomato^+^ microglia (arrowheads) and tdTomato expression in a large blood vessel (open arrowhead). **T-V**, Confocal micrograph showing tdTomato expression in cortical microglia and cells of the pia (open arrowheads). **W-Z**, Confocal micrograph showing CD163-(white) and IBA1-expressing (green) macrophages. The open arrowhead indicates a tdTomato^−^CD163^+^IBA1^+^ perivascular macrophage. Closed arrowheads parenchymal microglia. One of three independent experiments shown (N=3 mice).

The flow cytometry data revealed that recombination was largely, but not 100 percent specific. Furthermore, when examining the sections, we noticed some tdTomato fluorescence that did not appear to be associated with parenchymal microglia. To explore the nature of this ectopic fluorescence in this dosing paradigm, we imaged cells in the choroid plexus (Figure 5N-P) as well as cells in the cortex, which were immunostained for the endothelial cell marker CD31 (Figure 5Q-S). In the choroid plexus, there was very weak tdTomato fluorescence and most IBA1-positive macrophages did not express tdTomato (Figure 5N-P). In the cortex, high power confocal micrographs showed that the tdTomato and the CD31-immunolabeling signal were largely separate, with some tdTomato overlap with larger CD31-positive blood vessels (Figure 5Q-S). Furthermore, tdTomato fluorescence was present on the pia in IBA1-negative cells (Figure 5T-V) and absent from CD163-expressing perivascular macrophages (Figure 5W-Z). Together, these data indicate that the *Tmem119-CreERT2* line can be used to conditionally control gene expression in adult and early postnatal microglia with good specificity.

### Characterization of transgene expression in monocytes from Tmem119 knock-in mice

Monocytes infiltrate the brain and contribute to pathology in a number of brain disorders. Currently widely used transgenic microglia lines also target monocytes in addition to microglia and cannot readily distinguish the two cell types, thus hampering unequivocal attribution of microglia function in these disorders. To assess the utility of *Tmem119* knock-in mice for investigating microglia in models of CNS disorders, we isolated blood monocytes and examined transgene expression (Figure 6A). Using flow cytometry on white blood cells isolated from control or *Tmem119-EGFP^+/−^* mice, we did not detect any EGFP-expression (Figure 6B-C). To further determine whether *Tmem119-CreERT2^+/−^*; *Ai14*^+/−^ mice display recombination in monocytes, we administered tamoxifen and sacrificed the mice for analysis 7 days after the last dose (Figure 6D). Using flow cytometry on white blood cells, we found that about 3 percent of CD45^+^ cells expressed tdTomato (Figure 6E-F). This suggests that there is a low level of CreERT2 activity in monocytes of the *Tmem119-CreERT* mice. Together, these data indicate high applicability of *Tmem119-EGFP* mice and good applicability of *Tmem119-CreERT2* mice to the study of brain disorders with potential monocyte contribution.

**Figure 6.**
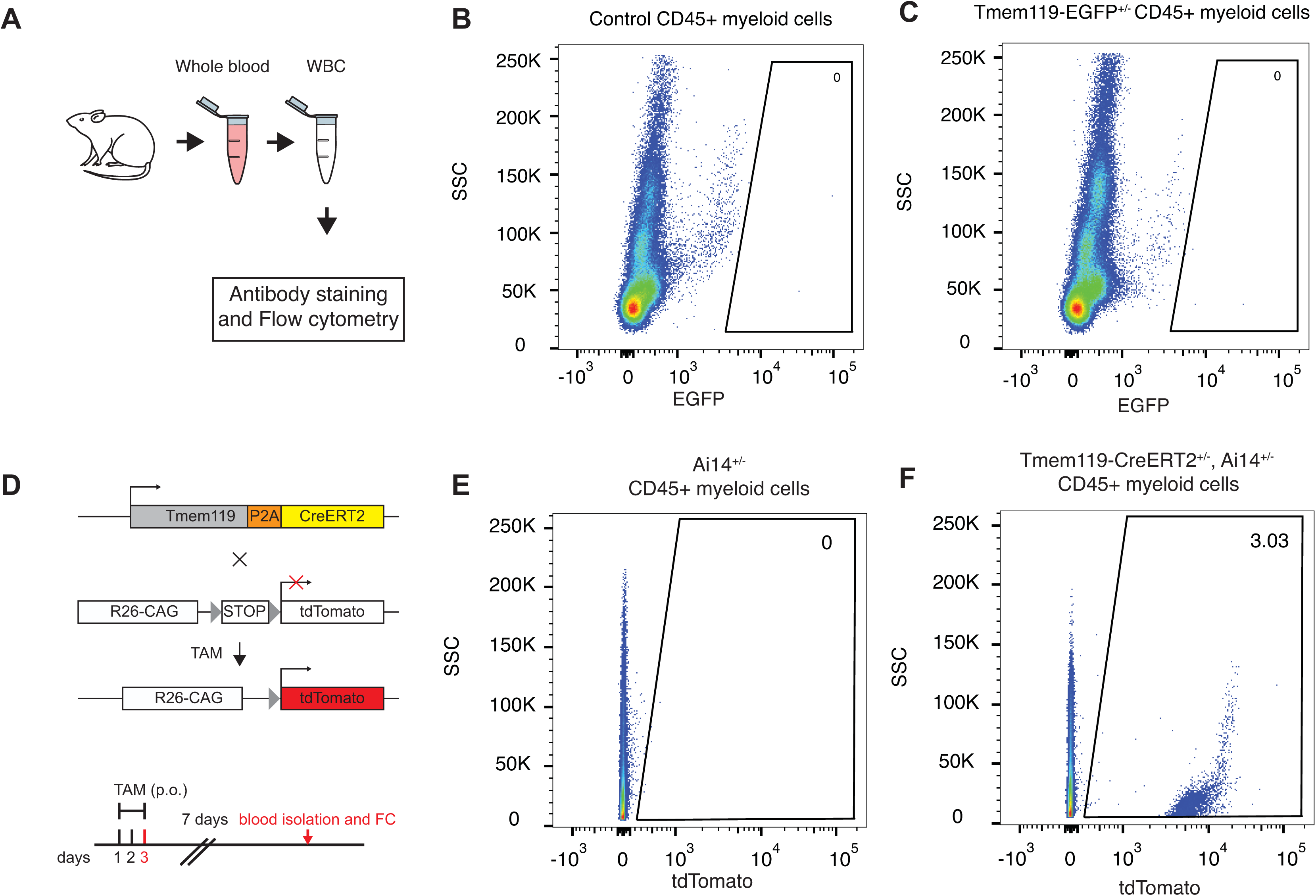
Blood monocyte profiling in Tmem119 knock-in mice. **A**, Experimental approach for flow cytometry of monocytes isolated from whole blood. **B and C**, Representative flow cytometry analysis of EGFP expression in pre-gated single live CD45^+^ cells in wildtype control (B) and *Tmem119-EGFP^+/−^* mice (C). One of three experiments is shown (N=3 mice). Numbers in or adjacent to outlined areas indicate percent cells in each gate. **D**, Adult mice of *Tmem119-CreERT2^+/−^; Ai14(RCL-tdT)-D^+/−^* mice received tamoxifen (TAM) per os (p.o) for 3 days. The mice were sacrificed 7 days after the last dose for flow cytometry (FC). **E and F**, Representative flow cytometry analysis of tdTomato expression in pre-gated single live CD45^+^ cells in *Ai14(RCL-tdT)-D^+/−^* (E) and *Tmem119-CreERT2^+/−^; Ai14(RCL-tdT)-D^+/−^* mice (F). One of two independent experiments is shown (N=3 mice).

### Statistical table

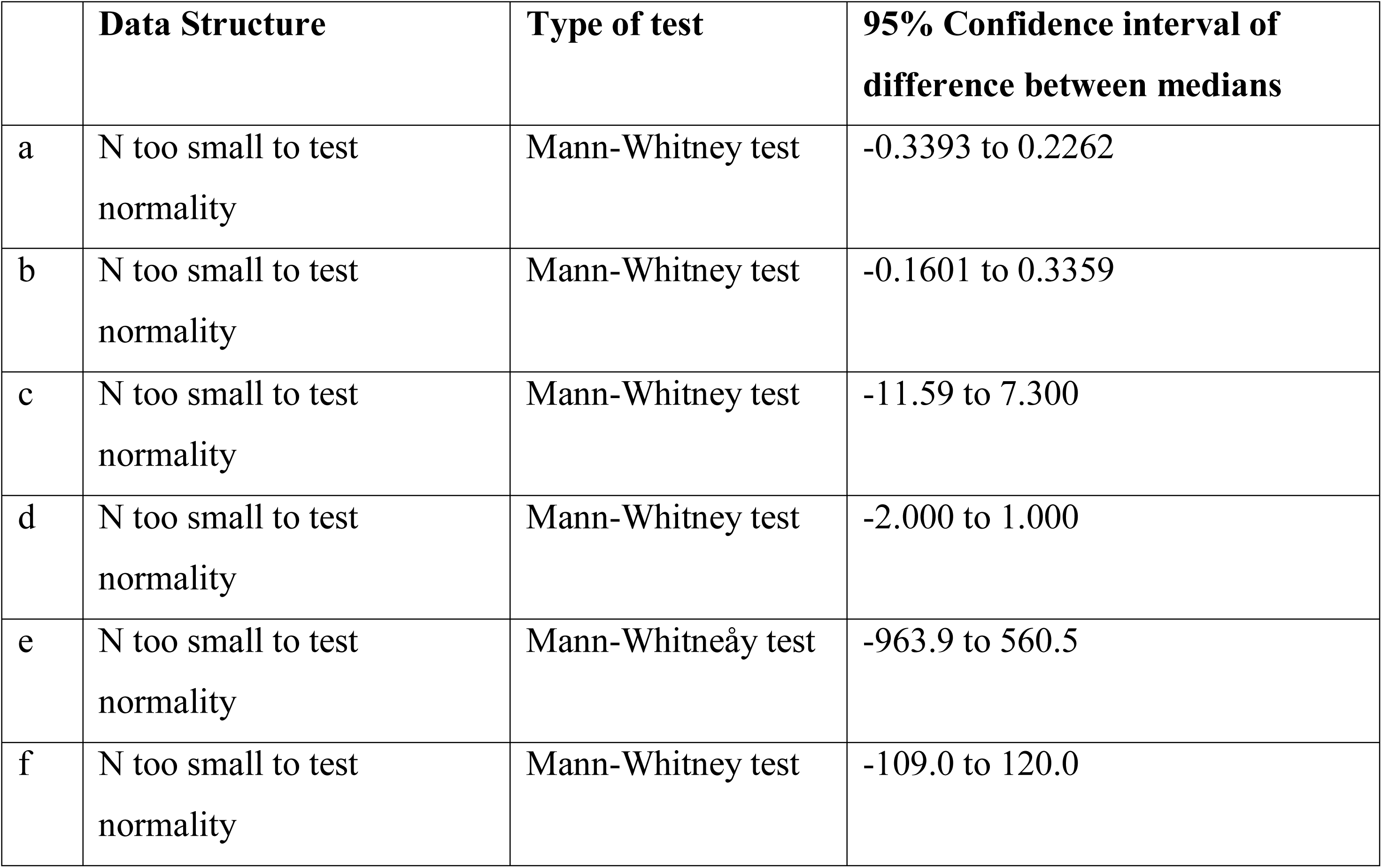

## Discussion

In this study, we addressed the critical need for transgenic mouse lines that specifically label or control microglia, but not other closely related cell types with overlapping signature gene expression (Bennett et al., 2016; Wieghofer and Prinz, 2016; Haimon et al., 2018). We targeted the recently identified microglia-specific *Tmem119* gene to knock in *EGFP* or *CreERT2*. Multiple analyses confirm correct insertion of the transgenes, intact expression of endogenous *Tmem119*, and normal basic morphology of microglia in the knock-in mice. Using immunofluorescence and flow cytometry approaches, we demonstrate that EGFP is expressed in microglia, but not other cells of the monophagocytic system defined anatomically and by signature protein expression. Flow cytometry further shows that EGFP is almost exclusively expressed in CD11b^+^CD45lo cells corresponding to microglia. We further provide evidence for early postnatal expression of EGFP and activity of CreERT2 in microglia upon tamoxifen administration. In addition, we show that blood monocytes do not express EGFP and display minimal activity of CreERT2. Together, our findings indicate proper generation of the two mouse lines, which are now readily available through a public repository (JAX#031823 for *Tmem119-EGFP* and JAX#031820 for *Tmem119-CreERT2*).

TMEM119 is a single transmembrane protein that is highly expressed in microglia in the brains of humans and mice (Bennett et al., 2016; Satoh et al., 2016). In this study, we opted to refrain from knocking *Tmem119* out through insertion of *EGFP* or *CreERT2* into the *Tmem119* ORF and instead chose to preserve expression by using a polycistronic knock-in approach with ribosomal skipping peptide P2A (Figure 1A). This was primarily motivated by the high expression levels of *Tmem119*, conservation of expression in microglia across species, and potential developmental regulation patterns, all of which suggest the importance of this gene. Disruption of important genes in microglia could lead to confounding haploinsufficiency phenotypes, as previously shown for other transgenic microglia lines (Lee et al., 2010; Rogers et al., 2011). Our data rule out such potential confounding effects by showing intact *Tmem119* mRNA and protein expression (Figure 1E-G), as well as normal basic morphology of microglia in the knock-in mice (Figure 1H-M).

Examination of EGFP expression and co-expression analysis with TMEM119, IBA1, and CD163 revealed that EGFP labels only parenchymal microglia (Figures 2 and 3). Identification of these cells was possible by direct observation of EGFP fluorescence as well as by analysis of immunostaining-enhanced tissue sections, supporting the conclusion that EGFP expression levels are robust. Flow cytometry analysis further showed expression of EGFP in virtually all CD11b^+^CD45lo microglia (Figure 2N) and a minor fraction of 6.9 percent (153/2212) of CD11b^+^CD45int/hi cells (Figure 3G-I). This reflects that *Tmem119* expression is highly selective, but not completely restricted to microglia, which has been observed in recent single-cell sequencing studies (Li et al., 2019). An interesting observation was the expression of EGFP at early postnatal time points in microglia (P1, P3, and P6), but also endothelial cells of blood vessels at P1 (Figure 4). This is interesting for two reasons. First, it suggests that *Tmem119* is initially expressed in microglia and endothelial cells and only becomes a specific microglia marker during early postnatal development, presumably by selective up- and downregulation of expression in microglia and endothelial cells, respectively. Second, translational processing of *Tmem119* transcript appears to stand in opposition to a previous report showing that, while RNA sequencing detects *Tmem119* mRNA throughout development, immunostaining only identifies TMEM119 protein at later postnatal stages (Bennett et al., 2016). Although our data also do not directly show TMEM119 protein expression at early stages, ribosomal processing of the transcript is at least indirectly implied by the presence of EGFP protein, suggesting TMEM119 protein might be expressed but undetected due to expression level, protein stability, differential posttranslational modification, or methodological limitations in immunostaining and flow cytometry. Together, these data indicate that *Tmem119* knock-in mice may enable the study of microglia and endothelial cells until postnatal day (P1) and microglia specifically after P3 and later.

Our experiments using *Tmem119-CreERT2^+/−^* Ai14(RCL-tdT)^+/−^ mice show tdTomato expression in IBA1-expressing microglia, indicating that this line is suitable for Cre-dependent manipulation of genes and transgene expression in microglia without affecting other major neuronal or glial cell types in the CNS (Figure 5). In this particular experiment, we observed very high, dose-dependent recombination efficiencies close to 100 percent with a trend for slight reduction of the generally high specificity with increasing tamoxifen dosage (Figure 5K, L). The recombination efficiency observed across all doses compares well with other published models, such as the *Cx3cr1* mouse line (Goldmann et al., 2013), and might even permit shorter dosing paradigms dependent on the assay and relevant conditional allele. We further examined tissues other than parenchymal microglia and observed that recombination was largely yet not completely absent in choroid plexus macrophages (5N-P). This is consistent with reported endogenous TMEM119 expression and other reports in the literature, suggesting other brain macrophages may express microglia-specific genes at very low levels (Bennett et al., 2016; Goldmann et al., 2016; Mildner et al., 2017), which might be sufficient to cause occasional all-or-none recombination events. In addition, our data showed tdTomato expression in a few larger blood vessels and IBA1-negative cells in the pia (Figure 5Q-V). While we did not determine the precise cellular origin of this signal, our data from CD31-immunostaining exclude endothelial cells of the intracerebral microvasculature (Figure 5Q-S). Together with the spatial extent of tdTomato expression and recent single cell RNA sequencing data of vascular cells, our observations point to leptomeningeal cells and their projections that penetrate deep into the brain ensheathing large blood vessels and the choroid plexus stroma (Decimo et al., 2012, Vanlandewijck et al., 2018). Together, these data render the *Tmem119-CreERT2* line a highly useful tool for the conditional study of microglia, while also suggesting potential application for leptomeningeal cells.

Examination of EGFP and tdTomato expression in blood monocytes of *Tmem119-EGFP* and *Tmem119-CreERT2; Ai14* mice revealed complete absence and minimal expression, respectively (Figure 6). While, the lack of EGFP expression in blood monocytes of the *Tmem119-EGFP* knock-in mice was expected based on previous observations, tdTomato expression in 3% blood monocytes of the *Tmem119-CreERT2; Ai14* mice was rather surprising. Similar to other brain macrophages discussed above, blood monocytes may express *Tmem119* at very low, almost undetectable levels, which may nonetheless be sufficient to cause occasional all-or-none Cre recombination events. Together, these data render the *Tmem119* knock-in lines powerful tools to investigate microglia in health and disease without significant confounds from monocyte contribution.

During the past decade, available microglia lines have been instrumental in advancing our understanding of microglia. These lines remain powerful tools to study microglia function. At the same time, it is widely acknowledged that ontogenetic differences, as well as differences in localization and microenvironment between parenchymal microglia and their closely related cells from the monophagocytic system warrant more discerning observation and manipulation (Bennett et al., 2016; Wieghofer and Prinz, 2016; Haimon et al., 2018). Recently developed antibodies for TMEM119 immunostaining are one tool that has been adopted quickly to identify parenchymal microglia. As TMEM119 is a transmembrane protein localizing to processes, immunostaining for this protein is particularly powerful to study morphology of microglia. To identify discrete microglia, however, relatively diffuse staining patterns of membrane-bound TMEM119 present a challenge (Figure 2B). In this case, the more discretely identifiable cytosolic distribution of EGFP in the *Tmem119-EGFP* line might be of great value. In addition, the robustness of the EGFP signal in the *Tmem119-EGFP* line may enable cell sorting without the need for antibodies, possibly even at early developmental stages before the onset of TMEM119 appearance on the membrane, as well as *in vivo* imaging approaches. Furthermore, the absence of EGFP fluorescence in blood monocytes renders this line uniquely useful for the study of microglia in brain disorders that involve monocyte infiltration. The *Tmem119-CreERT2* line will significantly facilitate conditional control of gene expression selectively in microglia, especially in cases when monocytes, but not leptomeningeal cells, could likely confound interpretations. Compared to constitutive Cre lines, which cumulatively recombine throughout development and thus deny temporal control, the *CreERT2* line will allow precise control of gene expression to examine the role of microglia in temporally restricted developmental processes. In this capacity, dose-dependency of CREERT2 and potentially resulting mosaic expression of recombined alleles using low doses may allow the study of candidate genes in adjacent microglia populations sharing the same microenvironment. Beyond the application of *Tmem119*-knockin and other currently available mouse lines, genetic approaches such as targeting other microglia-signature genes, engineering highly specific artificial promoters and regulatory regions, and using intersectional split-Cre approaches hold great promise to produce even more specific tools to study microglia in the future.

